# Genetic drivers shape TREM2-dependent myeloid responses and glioblastoma progression

**DOI:** 10.64898/2026.10.26.759860

**Authors:** Mekenzie M. Peshoff, Tanvi Joshi, Darwin D’Souza, Dimitrios Kleidonas, Mastura Akter, Alan Cieslukowski, Jordy Sepulveda, Jiaqian Wu, Dolores Hambardzumyan, Long-Jun Wu

## Abstract

Glioblastoma (GBM) is characterized by a heterogeneous, myeloid-enriched, and highly immunosuppressive microenvironment. TREM2 regulates microglial and macrophage functions, but its role in GBM remains unclear. We investigated whether genetic drivers determine the effects of TREM2 using *de novo* and orthotopic transplantable murine models of *Nf1*-silenced/Ras-driven and PDGFB-driven GBM. To this end, we combined immune profiling, functional assays, secretome analysis, and single-cell RNA sequencing with bone marrow chimeras and conditional *Trem2* deletion strategies. *Trem2* deficiency prolonged survival and reduced tumor burden in *Nf1*-silenced/Ras-driven GBM, whereas it increased tumor burden without affecting survival in PDGFB-driven GBM. These different outcomes were accompanied by distinct changes in myeloid composition and soluble mediators. TREM2 was required for bone marrow-derived macrophage migration and microglial process extension towards Ras-derived, but not PDGFB-derived, tumor-conditioned medium. In contrast, both models showed impaired microglial efferocytosis. Single-cell RNA sequencing analysis of *Nf1*-silenced GBM showed that *Trem2* loss shifted myeloid cells from disease-associated, pro-inflammatory states to a more homeostatic state. Bone marrow chimeras supported a tumor-promoting role for TREM2 in microglia, rather than infiltrating myeloid cells in Ras-driven GBM. In addition, conditional deletion of *Trem2* before tumor implantation prolonged survival, whereas deletion after tumor establishment did not. These findings identify genetic context and timing of TREM2 activity as determinants of its effects on GBM progression, highlighting driver-dependent myeloid responses as considerations for developing TREM2-directed therapies in glioblastoma.

## Introduction

Gliomas are the most common malignant primary brain tumors in adults.^1^ Of these, glioblastoma (GBM), isocitrate dehydrogenase (IDH)-wildtype (grade IV), is the most lethal, with a median survival of only 12-18 months.^2^ Standard-of-care treatment includes maximal safe surgical resection, followed by radiotherapy and concomitant and maintenance temozolomide^3^, yet only 6.8% of patients remain alive 5 years after diagnosis.^4^ Although immune checkpoint blockade has transformed the treatment of several solid tumors, its efficacy in GBM remains limited.^5,6^ Chimeric antigen receptor (CAR) T-cell therapy has shown promising preliminary anti-tumor activity in early-phase trials in some patients with recurrent GBM; however, response durability and survival benefit remain to be established.^7,8^ Recent clinical profiling linked an immunosuppressive myeloid scavenger program to CAR T-cell refractory recurrent GBM, suggesting that myeloid cells may limit treatment efficacy.^9^ Thus, the heterogeneous, myeloid-dominated microenvironment of GBM presents both a barrier to effective immunotherapy and an opportunity to improve treatment responses by targeting immunosuppressive myeloid cells.^10^ Understanding myeloid heterogeneity is essential to identify targets that improve immunotherapy across tumors versus those that work only in specific tumor types.

Approximately 30-40% of the GBM mass is attributable to tumor-associated myeloid cells (TAMs), which are primarily brain-resident microglia and monocytes, monocyte-derived macrophages (MDMs), and neutrophils that infiltrate from the periphery.^10,11^ While some TAMs are responsible for anti-tumor immune functions such as phagocytosis and antigen presentation^12,13^, these beneficial roles are largely subverted to support an immunosuppressive microenvironment.^14,15^ Functional phenotypes and relative proportions of TAM subsets can be shaped by GBM-intrinsic characteristics, including oncogenic driver mutations.^16–18^ Early molecular classifications of GBM identified proneural (PN), classical (CL), and mesenchymal (MES) subtypes, associated with key genetic alterations, including loss of the tumor suppressor Neurofibromin-1 (*NF1*) driving aberrant Ras activity in MES, amplification of Platelet-derived growth factor A (*PDGFRA*) in PN, and truncated *EGFRvIII* resulting in constitutive activation of epidermal growth factor receptor (EGFR) signaling in CL gliomas.^19–22^ A growing body of work refined by single-cell RNA sequencing indicates myeloid compartments in genetically-engineered mouse models with *Nf1-*silencing, *PDGFB-*overexpression, and *EGFRvIII-*expression (*Nf1* mGBM, *PDGFB* mGBM, *EGFRvIII* mGBM) closely recapitulate characteristics of human GBM.^17,23^ Thus, these categories and representative animal models provide an important framework for understanding how myeloid cell properties are shaped by glioma-intrinsic factors. Defining these relationships may reveal shared myeloid targets across GBM subtypes and distinguish them from those that require tumor-specific therapeutic strategies.

Triggering receptor expressed on myeloid cells 2 (TREM2) is a signaling hub for key cellular processes in microglia and macrophages, including metabolism, phagocytosis, and survival.^24,25^ Induction of TREM2 enables microglia to transition into a disease-associated phenotype during neurodegeneration, which is protective in early stages of Alzheimer’s disease and TDP-43 related proteinopathy.^26–28^ Due to its abundant expression on TAMs, TREM2 has also been studied in a variety of peripheral cancers, where it has largely been attributed to immunosuppression^29,30^; however, it can also be protective depending on tumor type.^31–33^ Because of its relevance in neurological disorders and cancer, we and others have investigated the role of TREM2 in gliomas with divergent results. While some reports have shown that *Trem2* deletion extends survival in some murine models of GBM^34,35^, others did not observe this effect and in some models, observed increased tumor burden in *Trem2^-/-^*groups.^36–38^ However, the basis for these divergent results remained unclear.^39^ Together with the substantial heterogeneity of TAMs, these findings led us to hypothesize that TREM2 does not have a uniform pro- or anti-tumor function across GBM subtypes, but instead regulates myeloid cell responses to distinct microenvironments shaped by tumor-intrinsic genetic alterations and transcriptional programs.

To test this hypothesis, we used *de novo* and orthotopic transplant murine GBM models with distinct oncogenic drivers. *Trem2* deficiency improved outcomes in Ras-driven (orthotopic transplant murine GBM, Ras oGBM) and *Nf1* mGBM but not *PDGFB* mGBM and PDGFB-driven orthotopic transplant murine model (PDGFB oGBM), with corresponding differences in the tumor immune microenvironment (TME). Functional assays and proteomics identified TREM2-dependent myeloid migration response to factors secreted by Ras-driven, but not PDGFB-driven, GBM cells. Single-cell RNA sequencing revealed TREM2-dependent changes to microglia and MDM subsets in *Nf1* mGBM. Conditional *Trem2* deletion and bone marrow transplantation further showed that disease stage and cell type influence TREM2 function in GBM. Together, these results help explain divergent results across models and show that TREM2 regulates myeloid cell responses to distinct GBM-derived signals in a context-dependent manner.

## Materials and methods

### Animal models

#### Husbandry and approval

All mice were maintained under standard housing conditions at the University of Texas Health Science Center (UTHealth), the Icahn School of Medicine at Mount Sinai, and the Perelman School of Medicine, University of Pennsylvania. Mice were given access to food and water *ad libitum* under a 12-hour light/dark cycle. All procedures were approved by the UTHealth Animal Welfare Committee (Protocol #AWC-24-0092), the IACUC (Institutional Animal Care and Use Committee) at the Icahn School of Medicine at Mount Sinai (Protocol #2019-00619 and #2014-0229), and the Perelman School of Medicine, University of Pennsylvania (Protocol #807737). Genotyping of all mice was done *via* Transnetyx (Cordova, TN).

#### Mouse strains

Wildtype C57BL/6J mice (WT) were purchased from Jackson Laboratory (JAX #000664) and maintained in-house. *Trem2* general knockout mice (*Trem2^-/-^*) were a gift from Dr. Marco Colonna (Washington University). To generate *CX3CR1^GFP/+^Trem2^-/-^* mice, *Trem2^-/-^*mice were crossed to *CX3CR1^GFP^* knock-in/knock-out (JAX #005582). *Trem2^flox/flox^* (JAX #029853) mice were crossed to *CX3CR1^CreER^* knock-in/knock-out (JAX #021160) to generate the *Trem2* conditional knockout line (*Trem2^flox/flox^CX3CR1^CreER/+^*). PepBoy (CD45.1 WT, JAX #002014) mice were used for bone marrow transplant experiments. *Ntv-a; Trem2^-/-^* mice were generated by crossing *Trem2* knockout mice from Jackson Laboratory (JAX #027197) with Nestin-*Tv-a^+^* mice.

#### Orthotopic GBM models

Male and female 6–12-week-old mice were maintained on 0.2 mg/ml ibuprofen in the drinking water for 72 hours pre- and post-operation. Immediately prior to surgery, mice were given 2.5 mg/kg subcutaneous 0.25% bupivacaine at the surgical site for additional analgesia. Mice were placed into a stereotaxic frame under isoflurane (1.5%) anesthesia. Under sterile conditions, a 0.5 cm longitudinal incision was made in the scalp, and a small burr hole was drilled in the skull ±1.5 mm AP and ±1.5 mm ML to the bregma. A 26-gauge 10 µL Hamilton syringe was lowered 3.5 mm into the brain and 2 µL of sterile PBS containing 1.0×10^5^ dnp53-HRasV12 or 5.0×10^5^ dnp53-PDGFB cells (gift from Dr. Q. Richard Lu) were injected into the dorsolateral striatum. The incision was closed with 3M VetBond tissue adhesive. Mice were monitored for recovery for the first 3 days after surgery. For survival studies, endpoint criteria included >20% weight loss, visible bump on the head, lethargy, or neurological sequelae including seizures. All mice were sacrificed at a humane endpoint.

#### Generation of *de novo* murine GBM

Chicken fibroblast cells (DF-1; ATCC CRL-3586) were grown at 39°C, 5% CO2 according to supplier instructions. Cells were transfected with RCAS-*hPDGFB*-*HA*, RCAS-*hPDGFA*-myc, RCAS-shRNA-*p53*-RFP, RCAS-shRNA-*Nf1,* and RCAS-shRNA-*Pten*-RFP using the Fugene 6 transfection kit (Promega, E5911). To generate tumors, approximately 40,000 transfected DF-1 cells were stereotactically delivered using a Hamilton syringe with a 30-gauge needle in 1 µl Neurobasal medium per mouse. Sex-matched, 6–12-week-old mice were used. A mixture of equal parts RCAS-*hPDGFB*-*HA*- and RCAS-shRNA-*p53*-RFP-transfected cells was injected to generate *PDGFB*-driven GBM in the right frontal striatum (coordinates: AP −1.5 mm and right −0.5 mm from bregma; depth −1.5 mm from the dura surface). Similarly, a mixture of equal parts RCAS-*hPDGFA*-myc-, RCAS-shRNA-*p53*-RFP-, RCAS-shRNA-*Nf1*- and RCAS-shRNA-*Pten*-RFP-transfected cells was injected to generate *Nf1*-silenced GBM (coordinates: AP −0.0 mm and right −0.5 mm from bregma; depth −1.5 mm from the dura surface). For survival analyses, mice were monitored for signs of end-stage tumor burden such as >20% weight loss, visible bump on head, lethargy, and seizures. All mice were sacrificed at a humane endpoint.

#### Bone marrow transplantation

For bone marrow chimeric mice, recipient 6–12-week-old *Trem2^-/-^*or PepBoy mice were maintained on prophylactic antibiotics (95 mg/kg sulfamethoxazole/trimethoprim provided in drinking water) starting one week prior to irradiation. Recipient mice were anesthetized using isoflurane, placed into an adapted imaging manifold with a lead overhang (Somni Scientific EquaFlow5) and lethally irradiated (two 5.5 Gy doses four hours apart at a rate of 1.22-1.28 Gy/min) in an X-ray irradiator (MultiRad350). Donor mice were sacrificed, then femurs and tibia were collected and flushed with sterile 1X PBS. The resulting bone marrow cells were resuspended in PBS at a volume of 100 µL/mouse, then injected retro-orbitally into recipient mice at a 1:2 donor to recipient ratio. After 6-8 weeks of reconstitution, chimeric mice were implanted with orthotopic GBM as described.

#### *Ex vivo* two-photon imaging

*CX3CR1^GFP/+^* and *CX3CR1^GFP/+^Trem2^-/-^* mice aged 4-5 weeks were deeply anesthetized and decapitated. Brains were placed in ice-cold oxygenated artificial cerebrospinal fluid (ACSF) bubbled continuously with 95% O_2_ and 5% CO_2_. ACSF contained the following components in mM: 126 NaCl, 26 NaHCO_3_, 2.5 KCl, 1.25 NaH_2_PO_4_, 2 CaCl_2_, 2 MgCl_2_, and 10 glucose, with sucrose added to adjust the osmolarity to 300-320 mOsm. Coronal slices were cut at 300 um thickness on a vibrating blade microtome (Leica #VT1000S), then allowed to recover for 30 minutes in oxygenated ACSF before imaging. Slice imaging was performed at room temperature in a perfusion chamber continuously supplied with oxygenated ACSF at a flow rate of approximately 2 mL/min. Microglia were imaged 50-120 µm below the slice surface using a two-photon imaging system (Scientifica) equipped with a tunable laser (Chameleon Vision II; Coherent). A glass pipette containing TCM or control medium with 1:100 Alexa Fluor 594 was lowered next to microglia using a micromanipulator. Laser wavelength was tuned to 940 nm to image GFP^+^ microglia and locate the Alexa Fluor 594-containing glass pipette simultaneously. The pipette tip was positioned with care taken to avoid direct contact with microglial somata. Contents of the pipette were allowed to passively diffuse. Imaging was performed using a 20X water-immersion lens, and time-lapse images were acquired at 1 Hz with a resolution of 1024 x 1024 pixels. Laser power was maintained at 30-40 mW under the objective. To quantify microglial response to TCM, we measured microglial processes entering from the outer area (R_o_(0), 70 µm diameter) into the inner area (R_i_(t), 35 µm diameter) around the pipette as previously described.^40,41^ The microglial response at any given time is calculated as R(t)=(R_i_(t)-R_i_(0))/R_o_(0).

#### Immunohistochemistry and immunofluorescence

For immunofluorescence staining of cryosectioned brains, tumor-bearing mice were perfused intracardially with ice-cold PBS, followed by 10% formalin. Brains were collected, incubated in 10% formalin for at least 24 hours, and then transferred to a 30% sucrose solution at 4°C. After 3-4 days, brains were embedded in OCT, sectioned coronally on a cryostat at 30 µm, and mounted on charged slides. Slides were washed in PBS, then antigen retrieval was performed in pH 6.0 citrate buffer in a pressure chamber at approximately 90°C for 20 minutes. After slides were cooled to room temperature (RT) and washed in PBS, nonspecific staining was blocked at RT using 4% BSA in PBS containing 0.04% Triton-X for 1 hour. Primary antibody (Table 1) was added to the slide and incubated overnight at 4°C. The following day, slides were washed with PBS and incubated in secondary antibody for 1 hour RT. Slides were mounted on coverslips using mounting medium containing DAPI and images were analyzed using ImageJ.

For immunofluorescence staining of formalin-fixed, paraffin-embedded (FFPE) brains, slides were deparaffinized by passing through Neo-Clear and decreasing grades of ethanol followed by antigen retrieval in a boiling solution of citrate- or Tris-based buffer at pH 6 or 9 (Vector Labs, H-3300). The sections were allowed to cool on ice and then subjected to permeabilization with 0.3% Triton X-100 detergent solution in PBS. Blocking was performed for 1 hour at RT (or overnight at 4°C) using 0.1% Triton X-100 containing 2% BSA and 5% normal donkey serum (Jackson ImmunoResearch). Primary antibodies (Table 1) were diluted in PBS and incubated for 1 hour at RT or overnight at 4°C. Slides were incubated in secondary antibodies for 1 hour at RT, followed by counterstaining with DAPI and mounting in 70% glycerol solution. All images were taken on the FLUOVIEW FV4000 confocal microscope (Evident Scientific), keeping laser and software settings consistent across samples. To quantify GFAP-positive area, we captured at least 5 representative images per sample at 20X magnification from different tumor regions and averaged the values. Images were analyzed using the Threshold function in ImageJ (FIJI).

For immunohistochemical staining of FFPE brains, 5 μm sections and the Leica Bond RX automated slide stainer were used. After staining, slides were dehydrated by passing through increasing grades of alcohol and Neo-Clear, then mounted in Permount medium. Mounted slides were allowed to dry overnight and scanned with a whole slide scanner (Hamamatsu). The NDP.View2 software was used to capture multiple screenshots representative of the tumor, and quantification was done using the ImageJ software.

#### Flow cytometry

Tumor-bearing mice were perfused with ice-cold Ringer’s solution or PBS, after which brains were harvested on ice. Tumors were enzymatically dissociated using 0.5% collagenase D and DNase I to generate single-cell suspensions. Cells were enriched by density-gradient centrifugation in 30% Percoll supplemented with 10% FBS. Following removal of the supernatant, the cell pellet was treated with RBC Lysis Buffer for 1 minute, centrifuged, and resuspended in 100 μL of blocking buffer consisting of 2% FBS, 5% normal rat serum, 5% normal mouse serum, 5% normal rabbit serum, and 10 μg/mL anti-Fc receptor antibody (BioLegend, #101319) in FACS buffer. Samples were blocked on ice for 20 minutes and then stained with a primary antibody cocktail (Table 1) for 30 minutes on ice as previously described.^42^ Live/dead staining was performed with the Zombie UV viability dye (BioLegend, #423108) diluted in PBS-EDTA. Cells were washed in FACS buffer and fixed for 30 minutes at 4°C using fixation buffer (eBioscience, #5223-56 and #5123-43). Intracellular (IC) staining was performed for 30 minutes at RT using the indicated antibodies (Table 1) diluted in permeabilization buffer (Invitrogen #8333-56). Cells were then washed, resuspended in 300 μL of FACS buffer, and maintained at 4°C until acquisition. Samples were acquired on an Aurora Cytek spectral flow cytometer, and data were analyzed using FlowJo software.

### Cell culture

#### Glioma stem cell (GSC) culture

Base GSC media was made from DMEM/F12 (Gibco #11320033) containing 30% glucose (Sigma #G8270), sodium bicarbonate (Gibco #25-080-094), HEPES buffer (Gibco #15630106), L-glutamine (Stem Cell #07100), sodium pyruvate (Gibco #11-360-070), and 1% penicillin-streptomycin (Gibco #15-140-122). Dnp53-HRasV12 and dnp53-PDGFB GSCs were expanded in GSC growth medium, which was made from GSC base medium containing 20 ng/mL EGF (Stem Cell #78006.1), 20 ng/mL FGF (Stem Cell #78003), B27 supplement (Gibco #12587010), N2 supplement (Stem Cell #07152), and 0.2% heparin (Stem Cell #07980). The dnp53-HRasV12-tdTomato and dnp53-PDGFB-tdTomato lines were generated using lentiviral particles (Takara #0036VCT) and selected using 2 µg/mL puromycin. All cells were maintained at 5% CO_2_ at 37°C.

#### Primary microglia and bone marrow-derived macrophage (pBMDM) culture

Mouse pups aged P2-P6 were used for primary microglia culture. Brains were collected, and the cerebellum, meninges, and olfactory bulbs were removed. Tissue was mechanically and enzymatically dissociated with 2.5% trypsin (Gibco #15-090-046) and 10 mg/ml DNAse I (Millipore Sigma #DN25) for 12 minutes at 37°C. This suspension was filtered through a 100 µm cell strainer and cultured in DMEM/F12 containing 10% fetal bovine serum (FBS) and 1% penicillin-streptomycin in T-75 vented tissue culture flasks pre-coated with poly-D-lysine (Gibco #A3890401). Spent media was aspirated and replaced every 48 hours. After 12-15 days when the astrocyte layer was >80% confluent, microglia were detached using an incubator-shaker at 37°C at 200 rpm for 30-45 minutes. Microglia were replated in poly-D-lysine coated plates and used for experiments 24-48 hours later.

For primary bone marrow-derived macrophages (pBMDMs), femurs and tibia were collected and placed into a 600 µL microcentrifuge tube with a small hole placed into a 1.5 mL tube containing sterile PBS. The samples were centrifuged for 2 minutes at 1000 g, and the resulting pellet was resuspended in pBMDM media. pBMDM media was comprised of high-glucose DMEM-GlutaMAX (Gibco #10569044) containing 1% penicillin-streptomycin, 10% FBS, and 20% L929 supernatant as a source of M-CSF (Sigma #85011425). pBMDMs were grown for 4-7 days or until the 10 cm dish was >80% confluent. Cells were detached using TrypLE Express Enzyme (Gibco #12605010), neutralized in media, then plated for experiments.

#### Generation of conditioned medium

For tumor conditioned media (TCM), dnp53-HRasV12 and dnp53-PDGFB cells were cultured in GSC growth medium until sphere density or confluence reached 70%. Spent medium was removed and replaced with basal phenol red-free DMEM/F12 containing no additives (Gibco #21041025). After 24 hours, conditioned medium was collected and centrifuged for 10 minutes at 18,000 g at 4°C. The supernatant was filtered through a 0.2 µm polyethersulfone syringe filter and stored at -80 °C for future use, including LC/MS proteomics. For proteomics, 100 µg of protein was used for each sample.

#### Migration assay

For pBMDM migration toward GSCs, a modified Boyden transwell chamber (8.0 µm pore size, ThermoFisher #140629) was added to a 24-well plate. GSCs were added to the bottom well in co-culture medium (DMEM/F12 containing 2% FBS and 1% penicillin-streptomycin) and allowed to condition the medium overnight. In the top well, pBMDMs were added in co-culture medium and incubated at 37°C and 5% CO_2_ for 4 hours. Medium and non-migrated cells were removed from the top of the membrane using a cotton swab, then migrated cells on the bottom membrane were fixed in 10% formalin for 15 minutes at RT. Membranes were washed gently in 1X PBS, then stained with 0.2% crystal violet (Millipore Sigma #V5265). After a final wash in PBS, membranes were cut from the Boyden chamber using a razor blade, then mounted onto a coverslip for imaging. Analysis was performed in ImageJ using the threshold tool and percent of crystal violet positive area was recorded.

#### Scratch wound healing assay

100,000 pBMDM were added to a 12-well plate in pBMDM culture medium and incubated at 37°C until confluence reached approximately 80%. Half of the medium was replaced with TCM, and pBMDMs were incubated 24 hours. A small wound was etched through the center of the well using a 200 µL pipette tip. Images were acquired every 3 hours using a Nikon AR1 microscope and analyzed using the ImageJ scratch wound plugin. Time-lapse images were normalized to the area at 0 hours of the same well.

#### Protein collection from tumor lysates

Tumors were isolated and dissociated using a tissue homogenizer (ThermoFisher #12-141-361) in PBS containing 1% protease and phosphatase inhibitor cocktail (ThermoFisher #78442) and 1% Triton-X (ThermoFisher #AAA16046AE). The tumor lysate was incubated on ice for 20 minutes, then centrifuged 15,000 g for 20 minutes at 4°C. The resulting supernatant was collected and stored at -80°C, and protein concentration was measured using a Pierce BCA protein assay kit (ThermoFisher #23225). 200 µg protein was used for the Proteome Profiler Mouse XL Cytokine Array (R&D Systems #ARY028) according to manufacturer instructions. Luminex multiplex ELISA was performed according to manufacturer instructions (R&D Systems #LXSAMSM). Blots were visualized using a Bio-Rad ChemiDoc imaging system.

#### Magnetic cell sorting, RNA isolation, and rt-qPCR

For collection of RNA from myeloid cells, tissue was processed into single cells as previously described. Using the mouse CD11b^+^ positive selection sorting kit (Stem Cell #18970) according to manufacturer directions, myeloid cells were collected as a pellet and used for RNA isolation using the QIAGEN RNeasy Micro kit (#74004). RNA concentration was measured using NanoDrop, and >100 µg template RNA was used for cDNA synthesis. Using the iScript kit (Bio-Rad #1708890), cDNA was generated and stored at -20°C for rt-qPCR. The SYBR Green mastermix kit (Bio-Rad #1725271) was used for rt-qPCR on a Bio-Rad CFX96 system.

#### Single-cell RNA sequencing

scRNA-sequencing was done using the Chromium 3’ V3 platform (10X Genomics) as previously described and data analysis was also performed in accordance with previously published pipelines.^17,43–45^ Transcriptomic analysis of the astrocyte subcluster was performed according to the pipeline published by Heiland, et al.^46^ Astrocytes were scored for four transcriptional programs using predefined gene signatures and Seurat module scoring. Module scores were projected into a two-dimensional state space using the *Scpubr* package cellular-state framework^47^, with progenitor-like and mature programs defining one axis and alternative- and inflammatory-activation programs defining the orthogonal axis. Cells were assigned to their dominant transcriptional state according to their position in the resulting state space.

### Statistical analysis

Statistical analyses and number of biological replicates are described in figure legends. P-values were calculated by two-tailed t-test for normally-distributed data, Mann-Whitney test for data not distributed normally, one-way ANOVA for an independent variable with three or more groups, or two-way ANOVA for analyses containing two independent variables. Data are presented as mean ± standard deviation. All analyses were performed in GraphPad Prism v.11 or in R.

## Results

### *Trem2* deficiency improves survival in Ras-driven but not PDGFB-driven GBM-bearing mice

To understand if genetic drivers are associated with changes in *Trem2* expression, we first analyzed our published single-cell RNA sequencing dataset of *Nf1*, *PDGFB*, and *EGFRvIII* mGBM^17^ (Fig. S1A). Our previous work demonstrated that these mouse models provide an accurate framework for studying genotype-dependent changes in TAM subset proportions and phenotypes. *Trem2* was expressed predominantly in microglia and at lower expression levels in MDM regardless of the underlying driver mutations (Fig. S1B-C). These results are consistent with previous reports.^34–37^ Because TREM2 activity requires adaptor proteins DAP10 or DAP12 and acts within a larger group of correlated innate immune programs within the GBM TME, we then calculated a score for a *Trem2*-associated transcriptional module as previously described.^41^ There were no changes in expression of this *Trem2* pathway module across GBM models (Fig. S1D). Although module scores did not differ at the RNA level, we previously showed that *Nf1* mGBM has a higher abundance of microglia than *EGFRvIII* or *PDGFB* mGBM^16^, consistent with findings in the human MES GBM subtype.^48^

Next, to assess TREM2 function in *Nf1*/Ras-driven versus PDGFB-driven GBM, we used complementary *de novo* and orthotopic transplant murine GBM models in *Trem2* germline knockout mice. For *de novo* models (mGBM), we employed the RCAS/*tv-a* (replication-competent avian sarcoma-leukosis virus long terminal repeat with a splice acceptor/tumor virus A) system (Fig. 1A). By delivering defined genetic drivers to nestin-expressing neural stem/progenitor cells, this system allowed us to assess contributions of *Trem2* to formation of genetically distinct gliomas with the same cells-of-origin.^49^ We first confirmed *Trem2* transcript level reduction in *Ntv-a;Trem2^-/-^*mice by single-cell RNA sequencing and complete loss at the protein level by immunofluorescence (Fig. S2A&B). In *Nf1* mGBM, *Trem2* deficiency significantly increased median survival from 64 days to 82.5 days (Fig. 1B). In contrast, *Trem2* deficiency did not alter survival in mice bearing *PDGFB* mGBM (Fig. 1C).

**Figure 1:**
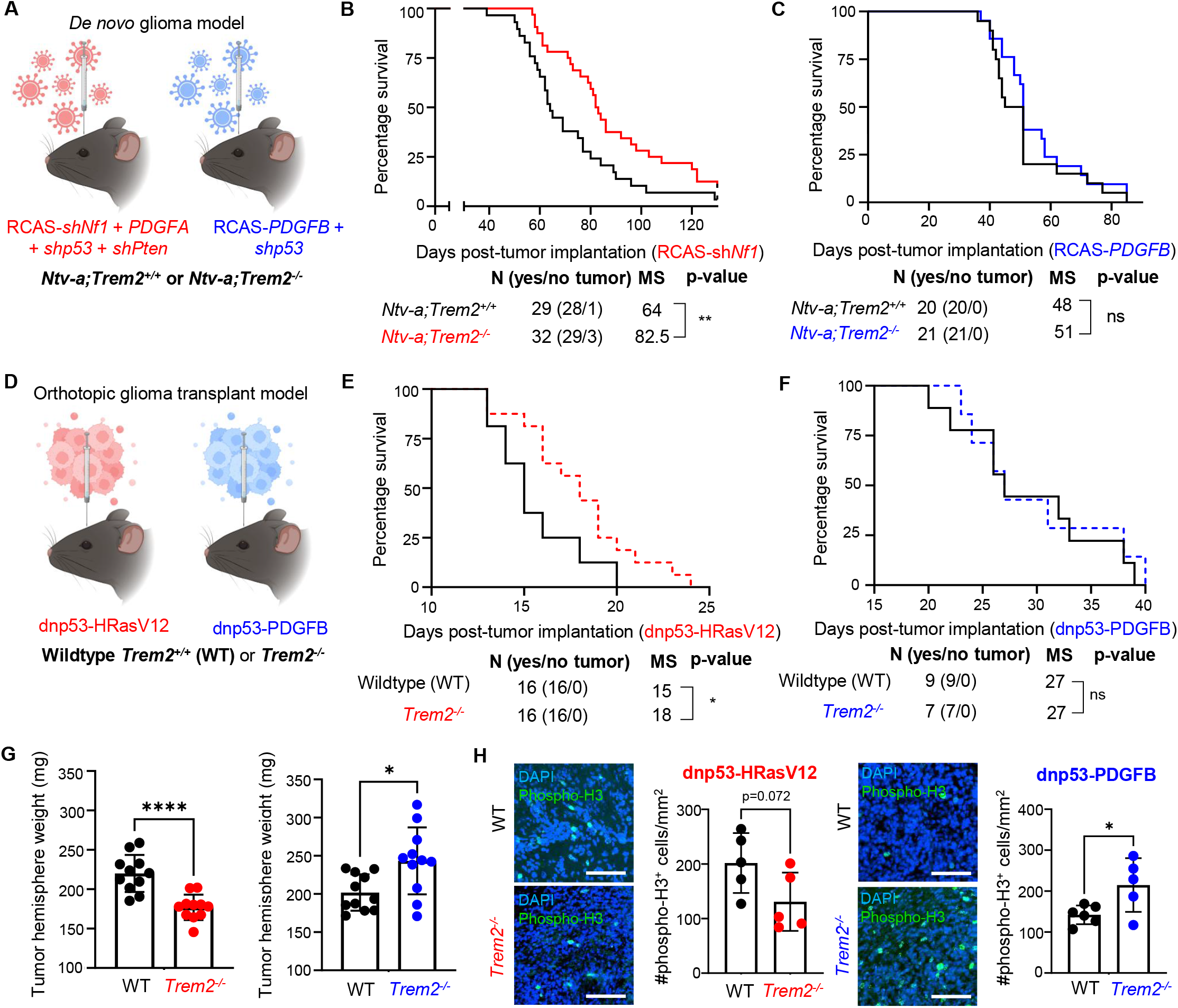
*Trem2* deficiency improves survival and tumor burden in Ras-driven, but not PDGFB-driven GBM. **(A)** Schematic of RCAS/*tv-a* model used to generate *de novo Nf1* (red) or *PDGFB* (blue) mGBM in *Ntv-a;Trem2^+/+^* and *Ntv-a;Trem2^-/-^*mice. Created in Biorender. **(B)** Kaplan-Meier curve comparing survival of *Ntv-a;Trem2^+/+^* (n=29) and *Ntv-a;Trem2^-/-^*(n=32) mice with *Nf1* mGBM; Gehan-Breslow-Wilcoxon test. **(C)** Kaplan-Meier curve comparing survival of *Ntv-a;Trem2^+/+^* (n=20) and *Ntv-a;Trem2^-/-^* (n=21) mice with PDGFB mGBM; Gehan-Breslow-Wilcoxon test. **(D)** Schematic of orthotopic Ras-driven (red) or PDGFB-driven (blue) oGBM in wildtype C57BL/6 or *Trem2^-/-^* mice. Created in Biorender. **(E)** Kaplan-Meier curve comparing survival of wild-type (n=16) and *Trem2^-/-^* (n=16) mice with Ras-driven oGBM; Gehan-Breslow-Wilcoxon test. **(F)** Kaplan-Meier curve comparing survival of wildtype (n=9) and *Trem2^-/-^* (n=7) mice with PDGFB-driven oGBM; Gehan-Breslow-Wilcoxon test. **(G)** Left: comparison of Ras-driven oGBM-bearing brain hemisphere weight in wild-type (n=11) and *Trem2^-/-^* (n=11) mice. Right: comparison of PDGFB-driven oGBM-bearing brain hemisphere weight in wildtype (n=11) and *Trem2^-/-^* (n=11) mice. Two-tailed, unpaired t-test. **(H)** Representative images and quantification of phospho-histone H3 (pHH3) in wild-type and *Trem2^-/-^* mice with Ras-driven and PDGFB-driven oGBM (n=5-6 per group). Two-tailed, unpaired t-test. Scale bar=100 µm. MS=median survival; *p<0.05, **p<0.01, ****p<0.0001.

We next examined whether the effects of *Trem2* deficiency also depended on the tumor’s oncogenic driver in orthotopic transplant models (oGBM), which involves the implantation of pre-differentiated glioma stem cells (GSCs; Fig. 1D) into wild-type (WT) or *Trem2^-/-^* mice. These two GSC lines were generated by retroviral vectors containing dominant-negative p53 and either an oncogenic Ras mutation (dnp53-HRasV12, Ras oGBM) or amplification of human PDGFB (dnp53-PDGFB, PDGFB oGBM) to complement the subtypes of the *de novo* models.^50^ In addition, the use of oGBM allowed us to match *in vivo* and *in vitro* phenotypes, as *Nf1* mGBM cells are unstable in culture.^51^ We found that *Trem2* deficiency extended median survival in mice bearing Ras oGBM (Fig. 1E) but did not alter survival in mice bearing PDGFB oGBM (Fig. 1F).

To assess the effect of *Trem2* on GBM establishment, we euthanized the mice at a set timepoint after orthotopic implantation (day 14 for Ras and day 21 for PDGFB oGBM). These timepoints were selected to capture comparable stages of disease progression in the two models.^50^ Compared to WT, *Trem2^-/-^* mice exhibited decreased tumor weight in Ras oGBM and increased tumor weight in PDGFB oGBM (Fig. 1G). The weights of the contralateral, non-tumor-bearing hemispheres did not differ significantly between genotypes, suggesting that the differences in tumor weight were not attributable to baseline differences in brain weight (Ras oGBM: average weight of 155.5 mg in WT and 154.7 mg in *Trem2^-/-^*; PDGFB oGBM: 161.8 mg in WT and 156.8 mg in *Trem2^-/-^*).

We next assessed phosphorylated histone H3 (pHH3) staining as a proxy for mitotic activity in these tumors. Consistent with tumor burden measurements, the number of pHH3-positive cells in *Trem2^-/-^* mice was dependent on genetic driver, with increased pHH3 staining in *Trem2^-/-^* PDGFB oGBM and decreased in *Trem2^-/-^* Ras oGBM compared to WT tumors (Fig. 1H). Together, these findings indicate that TREM2 affects GBM progression in a driver-dependent manner. *Trem2* deficiency increases tumor burden and mitotic activity in PDGFB oGBM without affecting survival, whereas it reduces tumor burden, decreases mitotic activity, and prolongs survival in Ras-driven gliomas.

### *Trem2* deficiency alters myeloid subsets and soluble mediators in a driver-dependent manner

We next asked whether the genetic driver-dependent effects of *Trem2* deficiency on glioma progression were also associated with differences in the TME. We performed immunohistochemical (IHC) staining for IBA1 in *Nf1* and *PDGFB* mGBM generated in *Ntv-a; Trem2^-/-^* mice (Fig. 2A-C). However, IBA1 staining alone cannot distinguish between microglia, MDM, monocytes, and some subsets of DCs; additionally, our previous work showed that *Nf1* and *PDGFB* mGBM contain varying proportions of myeloid populations.^23^ We therefore used multiplex spectral flow cytometry to distinguish individual cellular populations. Total myeloid cell abundance did not change with *Trem2* deficiency in either model; however, *Trem2* loss affected different subpopulations depending on the genetic driver. *Nf1* mGBM in *Ntv-a;Trem2^-/-^*mice displayed a reduction in microglia with no differences in infiltrating populations, whereas *PDGFB* mGBM generated in *Ntv-a;Trem2^-/-^* mice showed reduced BMDM (bone marrow-derived macrophages), monocytes, and neutrophils with no change in microglia (Fig. 2D&E). This observation is consistent with the enrichment of microglia in *Nf1* mGBM and monocyte lineage cells in *PDGFB* mGBM. Analysis of key phenotypic markers in microglia and MDM showed no differences in intensity or percentage of cells expressing MHC-II or PD-L1 between tumors generated in *Ntv-a; Trem2^+/+^* and *Ntv-a; Trem2^-/-^* mice for either model (Fig. S3A-D). Lymphoid populations also remained largely unchanged following *Trem2* deletion (Fig. S3E-I), although we did observe that CD4^+^ T-cells displayed increased TIM-3 expression in *PDGFB* mGBM generated in *Ntv-a; Trem2^-/-^* mice (Fig. S3J). This is consistent with previous reports indicating that TREM2 activates CD4-mediated immunity in GBM.^36^ The orthotopic transplant models showed similar driver-dependent changes in microglial abundance following *Trem2* deletion in Ras oGBM (Fig. S4A–D).

**Figure 2:**
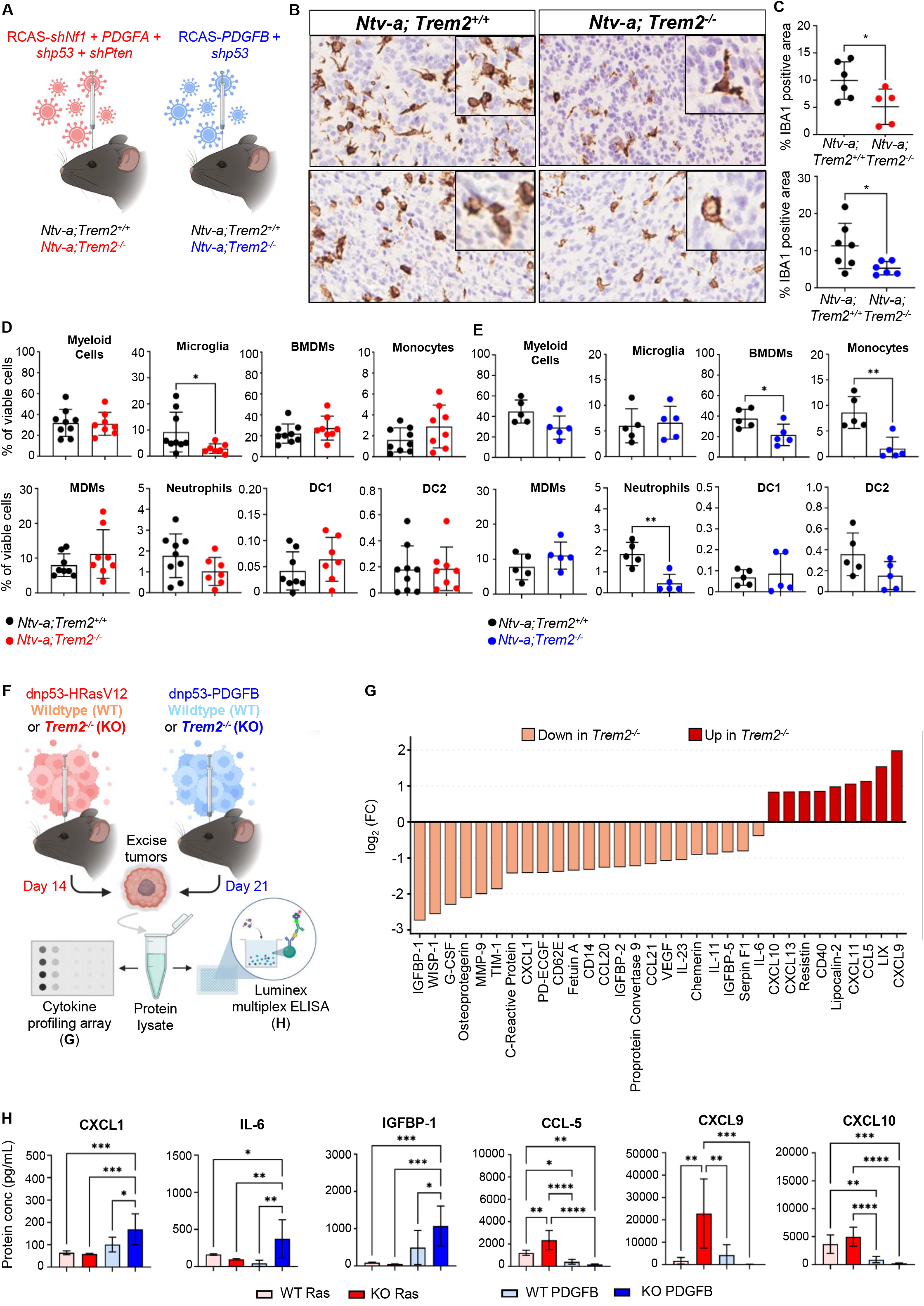
Immune profiling of *Trem2*-deficient and wild-type mice with Ras-driven or PDGFB-driven GBM. **(A)** Schematic of *de novo Nf1* (red) and *PDGFB* (blue) mGBM. Created in Biorender. **(B)** Representative images of IBA1 staining in *Ntv-a;Trem2^+/+^* and *Ntv-a;Trem2^-/-^ Nf1* (top) or *PDGFB* (bottom) mGBM. **(C)** Quantification of IBA1-positive area in *Ntv-a;Trem2^+/+^* and *Ntv-a;Trem2^-/-^ Nf1* (top) or *PDGFB* (bottom) mGBM, n=5-7 per group. **(D)** Proportions of myeloid cell populations in *Ntv-a;Trem2^+/+^* (n=9) and *Ntv-a;Trem2^-/-^* (n=8) *Nf1* mGBM by flow cytometry. Two-tailed, unpaired t-test. **(E)** Proportions of myeloid cell populations in *Ntv-a;Trem2^+/+^* (n=5) and *Ntv-a;Trem2^-/-^* (n=5) *PDGFB* mGBM by flow cytometry. Two-tailed, unpaired t-test. **(F)** Schematic showing cytokine profiling and Luminex ELISA workflow for wildtype and *Trem2^-/-^* oGBM. Created in Biorender. **(G)** Waterfall plot of fold changes for cytokines in Ras-driven wild-type and *Trem2^-/-^* oGBM lysates from cytokine profiling array. **(H)** ELISA of selected cytokines and chemokines in tumor lysates of wild-type and *Trem2^-/-^* Ras and PDGFB oGBM (n=6 for all groups). One-way ANOVA. *p<0.05, **\*\***p<0.01, **\*\*\***p<0.001.

Since TREM1 and TREM2 signal through a shared receptor, DAP12, and given that TREM1 is upregulated in GBM^52–54^, we also investigated the effects of *Trem1* deficiency in *PDGFB* and *Nf1* mGBM. Interestingly, while *Trem2*-deficient mice displayed reduced *Trem1*, *Trem1* deficiency was not sufficient to extend survival or change immune cell composition irrespective of driver mutations (Fig. S5A-I).

Next, we investigated whether *Trem2* deficiency changes soluble factors such as chemokines and cytokines in the TME. Using GSC culture models, we performed cytokine array profiling to screen for phenotypic changes (Fig. 2F). Several known GBM inflammatory mediators related to myeloid cell recruitment were increased in *Trem2^-/-^* Ras oGBM, including CCL5, LIX/CXCL5, and CXCL11 (Fig. 2G). Notably, *Trem2^-/-^* Ras oGBM also displayed elevated T-cell recruitment chemokines CXCL9 and CXCL10, although T-cell numbers remained largely unchanged at the tumor endpoint (Fig. S3E-F and Fig. 2G). In PDGFB oGBM, *Trem2* deficiency produced a distinct soluble-factor profile, with increased IL-6, CXCL1, MMP-9, and IGFBP-1 (Fig. S6A). The increase in these cytokines suggests enhanced signaling associated with immune regulation, matrix remodeling, immunosuppression, and angiogenesis, potentially contributing to the greater tumor burden and proliferation in *Trem2-*deficient PDGFB oGBM (Fig. 1G).^55–60^ To validate the cytokine array findings, we then performed Luminex multiplex ELISA using additional samples to quantify changes in cytokines of interest (Fig. 2H), which confirmed distinct changes to key soluble factors in *Trem2^-/-^* Ras versus PDGFB oGBM. Together, these findings indicate that *Trem2* deficiency remodels the tumor microenvironment in a driver-dependent manner, increasing inflammatory mediators in Ras-driven GBM while producing a mixed profile of immune-regulatory and tumor-associated factors in PDGFB-driven tumors.

### Myeloid-GBM communication displays driver-specific TREM2 dependence

To determine if contributions from the broader tumor microenvironment were required to elicit these genotype-dependent effects of *Trem2*, we employed *in vitro* and *ex vivo* functional assays to interrogate direct myeloid-glioma communication. We began by performing a migration assay with WT or *Trem2^-/-^* primary bone marrow-derived macrophages (pBMDMs) in the top well of a Boyden chamber, allowing them to migrate toward Ras GSCs, PDGFB GSCs, or a cell-free control (Fig. 3A). After 4 hours, the pBMDMs had migrated through the membrane and could be visualized upon crystal violet staining (Fig. 3B). WT pBMDMs exhibited increased migration toward Ras-driven GSCs compared to PDGFB-driven GSCs, whereas this difference was absent in *Trem2^-/-^*pBMDMs (Fig. 3C). pBMDMs can also coordinate their movement as a group through collective cytoskeletal arrangement, so we performed a scratch wound healing assay as an additional readout to evaluate the directed migration results from the Boyden assay. WT or *Trem2^-/-^* pBMDMs were incubated in control medium or tumor-conditioned medium (TCM) from Ras- or PDGFB-driven GSCs, then a scratch assay was performed (Fig. 3D). While *Trem2^-/-^* and WT pBMDMs displayed similar rates of wound healing after treatment with PDGFB TCM, the response after Ras TCM was markedly reduced in *Trem2^-/-^*(Fig. 3E&F).

**Figure 3:**
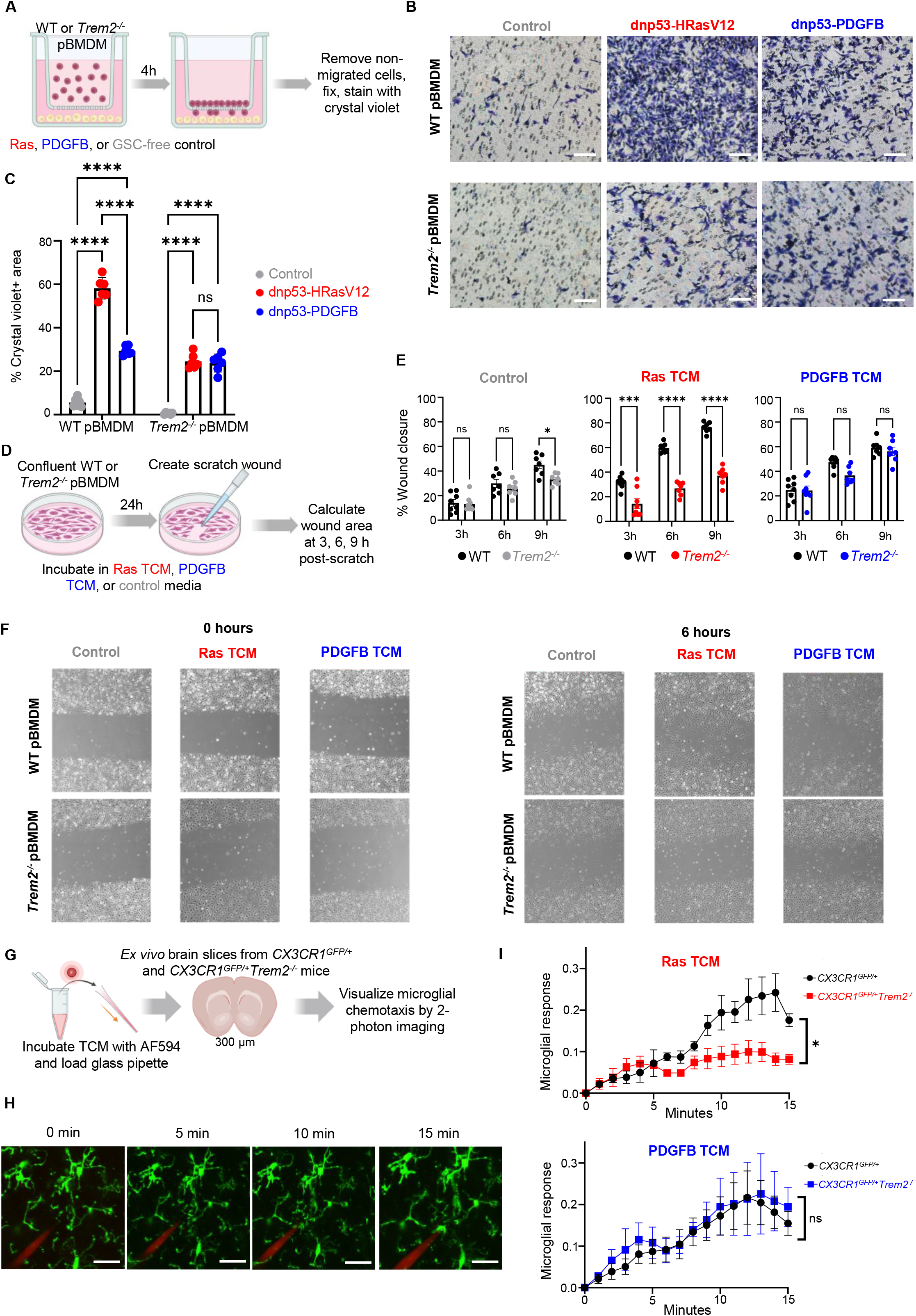
TAM migration toward Ras-secreted, but not PDGFB-secreted, factors is *Trem2*-dependent. **(A)** Schematic depicting migration assay of wild-type or *Trem2^-/-^*primary bone marrow-derived macrophages (pBMDM) toward Ras-driven or PDGFB-driven glioma stem cells (GSCs) or GSC-free control. Created in Biorender. **(B)** Representative images of wild-type or *Trem2^-/-^* pBMDM migration toward Ras, PDGFB, or control. Scale bar=100 µm. **(C)** Quantification of pBMDM migration via crystal violet area (n=6 per group). Two-way ANOVA. **(D)** Schematic of scratch wound healing assay with wild-type or *Trem2^-/-^* pBMDM incubated in Ras-driven or PDGFB-driven tumor-conditioned medium (TCM) or control medium. Created in Biorender. **(E)** Quantification of scratch wound healing assay for wild-type or *Trem2^-/-^* pBMDM incubated with control (gray), Ras-driven TCM (red), or PDGFB-driven TCM (blue). Two-way ANOVA, n=7-8 per group. **(F)** Representative images of scratch wound assay 0 hours and 6 hours post-scratch. **(G)** Schematic of *ex vivo* two-photon imaging of microglial chemotaxis toward TCM. Created in Biorender. **(H)** Representative images of *CX3CR1^GFP/+^* microglial chemotaxis toward TCM-containing pipette at 0, 5, 10, and 15 minutes. Scale bar=20 µm. **(I)** Quantification of microglia response to Ras- or PDGFB-driven TCM in *CX3CR1^GFP/+^* and *CX3CR1^GFP/+^Trem2^-/-^* brain slices (n=3 for all groups). Two-way ANOVA. *p<0.05, ***p<0.001, ****p<0.0001.

We previously showed that in adult GBM models, microglial somata are largely stationary^61^, and these cells rely on highly motile processes to extend and retract to survey their environment.^62,63^ Therefore, we chose a physiologically relevant model of microglial chemotaxis to assess how *Trem2* deficiency affects microglial response to Ras and PDGFB TCM. To visualize microglial dynamics, we used organotypic brain slices from *CX3CR1^GFP/+^* and *CX3CR1^GFP/+^Trem2^-/-^* mice and lowered a patch pipette filled with fluorescently labeled TCM onto the cortical surface (Fig. 3G). Microglia were able to extend their processes toward the TCM-containing pipette throughout a 15-minute window, allowing us to quantify their response using established methods (Fig. 3H).^40,41^ Consistent with pBMDM migration, *Trem2* deficiency impaired microglial chemotaxis toward Ras-derived, but not PDGFB-derived, TCM (Fig. 3I). Because we previously demonstrated the role of TREM2 in phagocytosis within the GBM microenvironment^37^, we also performed a microglial efferocytosis assay using apoptotic, tdTomato-expressing Ras and PDGFB GSCs as targets. *Trem2-*deficient microglia displayed impaired uptake of both Ras and PDGFB GSCs in phagocytosis assays (Fig. S7A-B), suggesting that impaired efferocytosis alone does not explain the driver-dependent effects of *Trem2* deficiency. These functional assays reveal genetic driver-dependent effects of TREM2 on pBMDM migration and microglial process extension toward glioma-derived TCM, with TREM2 dependence observed in the Ras model but not the PDGFB model.

### Ras- and PDGFB oGBM establish distinct, TREM2-relevant secretomes

To better understand the mediators secreted by Ras-driven and PDGFB-driven GBM tumors that may influence TREM2 function, we performed liquid chromatography/mass spectrometry (LC/MS) proteomic profiling of TCM from Ras and PDGFB GSCs (Fig. 4A). We detected 309 high-confidence proteins, including 166 unique to Ras TCM, 55 unique to PDGFB TCM, and 88 proteins secreted by both GSCs (Fig. 4B). Principal component analysis showed separated biological replicates from consecutive passages based on tumor genotype, with the first principal component accounting for 50.4% of the variance (Fig. 4C). Hierarchical clustering of the 50 most differentially abundant proteins likewise separated the Ras and PDGFB groups, highlighting distinct secretome profiles (Fig. 4D).

**Figure 4:**
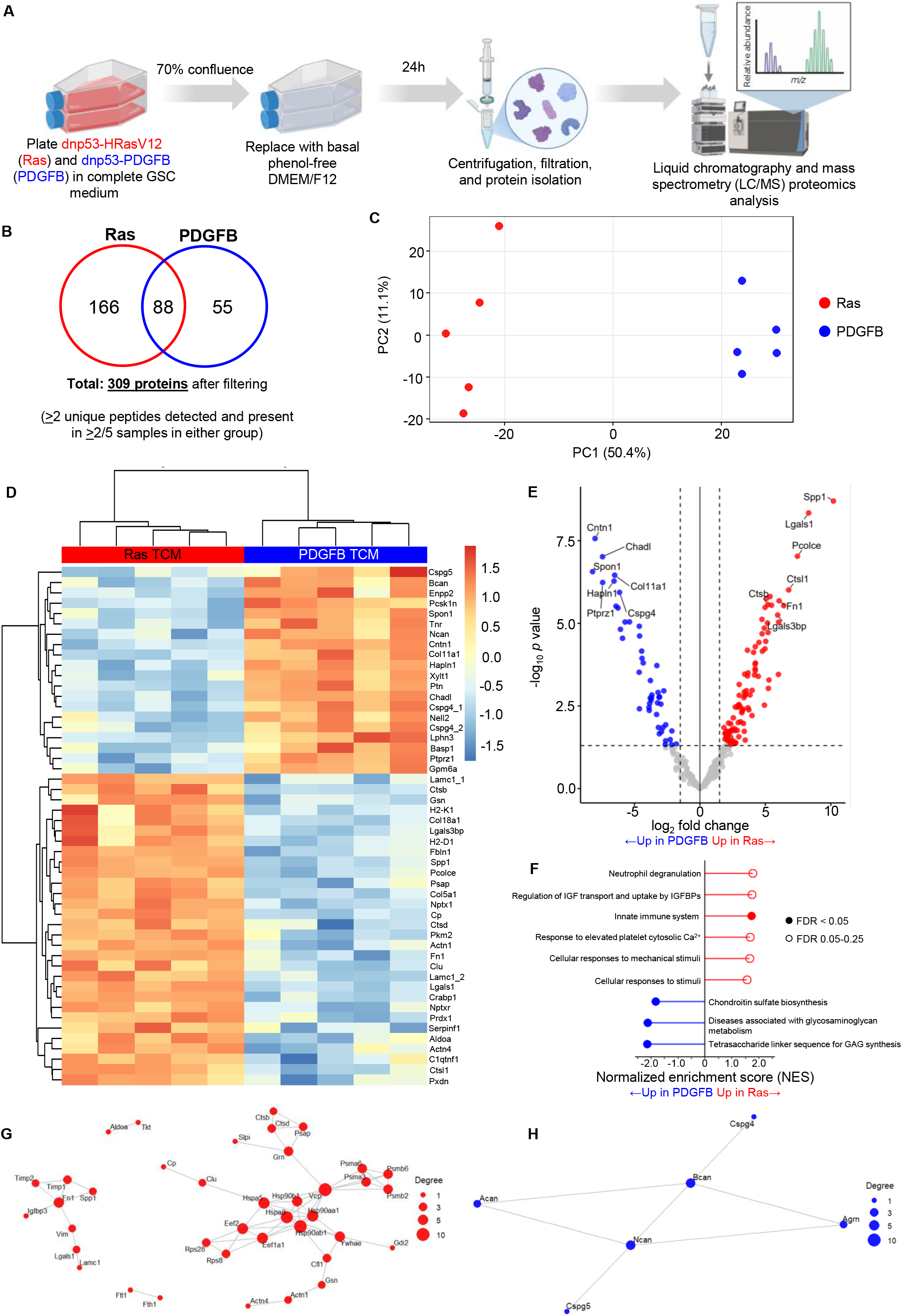
LC/MS profiling of tumor conditioned medium reveals distinct, *Trem2*-relevant secretomes of Ras-driven and PDGFB-driven GBM. **(A)** Schematic of tumor-conditioned medium (TCM) collection and profiling. Created in Biorender. **(B)** Venn diagram for number of proteins detected in Ras TCM (red), PDGFB TCM (blue), or both groups. **(C)** Principal component analysis (PCA) of Ras TCM and PDGFB TCM samples (n=5 for both groups). **(D)** Volcano plot of significant proteins (p<0.05 and |log2 fold change| >1.5) between Ras and PDGFB TCM. **(E)** Heatmap with hierarchical clustering for top 50 differentially abundant proteins between Ras and PDGFB. **(F)** GSEA analysis for top pathways enriched in Ras or PDGFB TCM. **(G)** Protein-protein interaction network in Ras TCM. (H) Protein-protein interaction network in PDGFB TCM.

Several proteins differed in abundance between Ras- and PDGFB-driven TCM (Fig. 4E). Ras-driven TCM displayed an enrichment of canonical disease-associated microglia factors including Spp1 (Osteopontin), Lgals1 (Galectin-1), and Lgals3bp (Galectin-3 binding protein)^64–68^, consistent with our previous reports of an Spp1-integrin axis in *Nf1* mGBM.^17^ Other proteins abundant in Ras TCM, such as Fn1 (fibronectin) and the cathepsins Ctsl1 and Ctsb, are indicative of chronic neuroinflammatory pathways.^69–72^ In contrast, PDGFB TCM represented a neurodevelopmental extracellular matrix program, including collagen and proteoglycan-associated proteins (i.e., Chadl, Ncan, Bcan, Hapln1).^73,74^ These distinct secretory profiles could provide different extracellular cues for TREM2-expressing myeloid cells, potentially contributing to context-dependent functions in Ras-driven and PDGFB-driven GBM.

GSEA pathway analysis further distinguished the Ras- and PDGFB-driven secretomes. While proteins high in Ras TCM were associated with cellular stimulus response and innate immune pathways, PDGFB-enriched proteins mapped to glycosaminoglycan (GAG) and chondroitin sulfate metabolism, consistent with their proneural-like identity (Fig. 4F). A protein-protein interaction analysis revealed a relatively interconnected network among the Ras-secreted factors (Fig. 4G), while PDGFB-enriched proteins form a smaller network centered around components of the neural extracellular matrix (Fig. 4H). Together, these results indicate that Ras and PDGFB GSCs establish unique extracellular niches. Notably, the Ras-enriched secretome supports disease-associated microglia polarization and cancer immunosuppression,^75–78^ providing candidate factors that could contribute to preferential engagement of TREM2-dependent myeloid cell migration.

### *Trem2* deficiency remodels myeloid cell transcriptional states in Ras-driven GBM

To assess transcriptional changes associated with *Trem2* deficiency, we sorted CD11b^+^ myeloid cells from wild-type and *Trem2^-/-^*Ras and PDGFB oGBM, then performed bulk RNA sequencing. In line with the lack of survival difference in PDGFB tumors, there were few changes in differential gene expression in *Trem2^-/-^* and WT sorted myeloid cells; however, there were abundant differences between *Trem2^-/-^*and WT Ras oGBM-derived myeloid cells and between different oGBM drivers within *Trem2^-/-^* or WT groups (Fig. S8A-D). Because of the survival benefit and pronounced *Trem2-*dependent functional responses, we then performed single-cell RNA sequencing (scRNA-seq) on endpoint *Nf1* mGBM from *Ntv-a;Trem2^+/+^* (n=3) and *Ntv-a;Trem2^-/-^* (n=3) mice. Following quality control, the dataset consisted of 51,310 cells and captured the major cell populations within the mGBM TME (Fig. 5A and Fig. S9A). Cell type composition analysis revealed a reduced fraction of tumor cells and an increased fraction of endothelial cells and astrocytes in the *Trem2*-deficient tumors (Fig. 5B-D and Fig. S9D). Cell cycle analysis did not reveal any changes between wild-type and *Trem2-*deficient tumors (Fig. 5E, Fig. S9B&C, Fig. S10B). We next performed additional immunohistochemical validation using pHH3 staining, resulting in no differences between the two groups (Fig. S13A&B). To further characterize the tumor compartment, we identified tumor cells based on *RCAS* expression (Fig. S10A). We next projected the four GBM cellular states from Neftel, *et al.*^79^ onto our dataset (Fig. 5F and Fig. S10C). Interestingly, we observed a difference in the AC-like tumor state, which was reduced in *Trem2-*deficient mice, along with a decrease in the mean module score for this state (Fig. 5G and Fig. S10D). Differential gene expression analysis of the tumor subcluster between the two genotypes revealed a downregulation of several genes associated with translation, ribosomal machinery and oxidative phosphorylation in the *Trem2*-deficient tumors (Fig. S10E).

**Figure 5:**
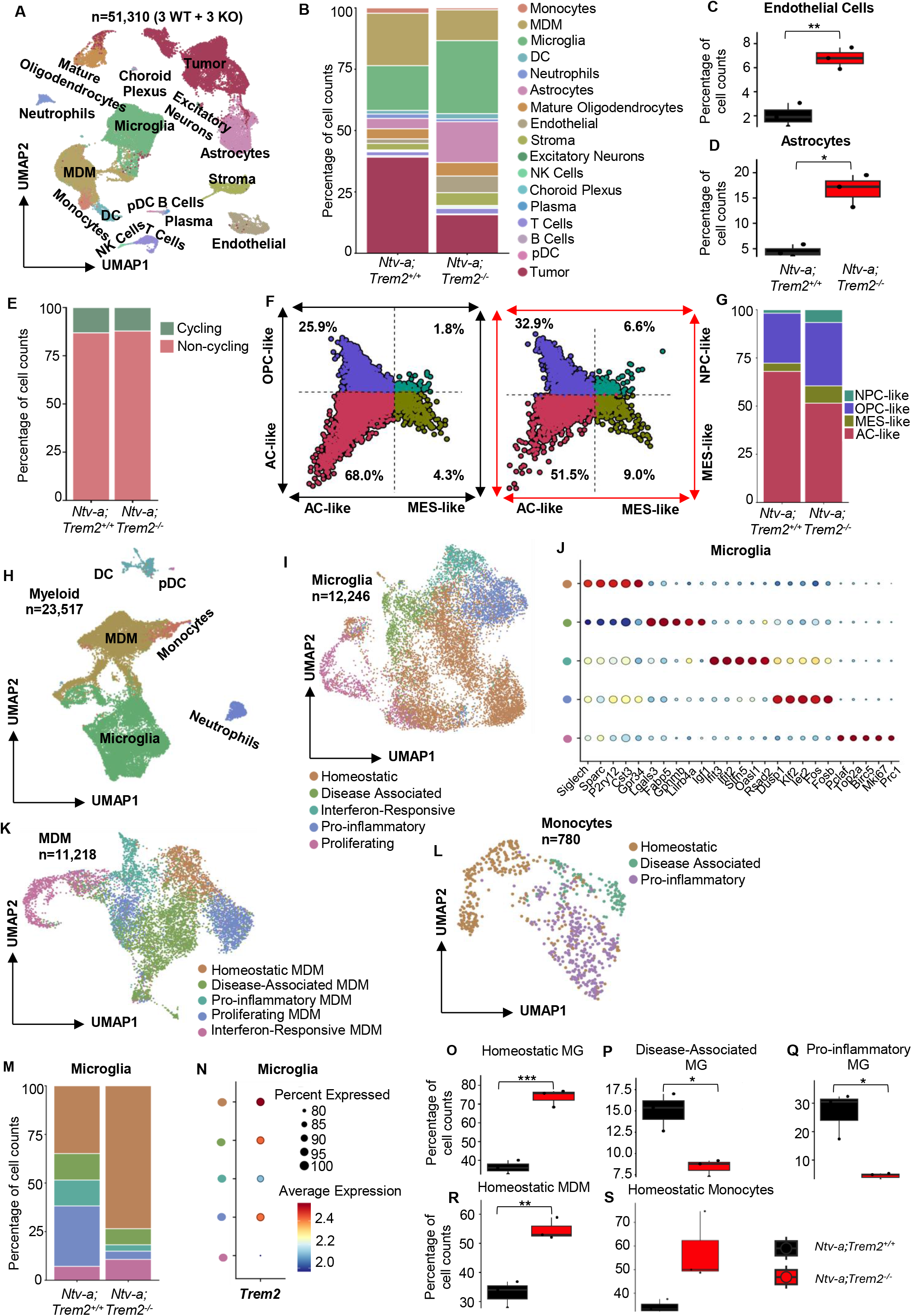
*Trem2* deficiency results in a reduction of disease-associated myeloid subpopulations in *Nf1* mGBM. **(A)** Uniform manifold approximation and projection (UMAP) dimensionality reduction of all cell populations within the scRNA-seq dataset. **(B)** Bar plot showing the proportion of each cell type from the UMAP compared between n=3 *Ntv-a;Trem2^+/+^* and n=3 *Ntv-a;Trem2^-/-^* Nf1 tumors. **(C and D)** Box plots showing the endothelial cell and astrocyte subpopulations respectively in *Ntv-a;Trem2^+/+^* (n=3) and *Ntv-a;Trem2^-/-^* (n=3) *Nf1* tumors as a percentage of all cells. Wilcoxon rank-sum test *p<0.05 **p<0.01. **(E)** Bar plots depicting the proportion of cycling and non-cycling cells from the UMAP compared between *Ntv-a;Trem2^+/+^*(n=3) and *Ntv-a;Trem2^-/-^* (n=3) *Nf1* tumors. **(F)** A two-dimensional quadrant scatter plot of the Neftel *et al* GBM cellular states projected onto *Ntv-a;Trem2^+/+^* (n=3) and *Ntv-a;Trem2^-/-^* (n=3) *Nf1* tumors. Astrocyte-like cells (AC) are depicted in pink, oligodendrocyte precursor cell-like cells (OPC-like) are depicted in purple, neural progenitor cell-like cells (NPC-like) are depicted in dark green and mesenchymal-like cells (MES-like) are depicted in light green. **(G)** Bar plot showing the proportion of each of the Neftel GBM cell states from (F) between *Ntv-a;Trem2^+/+^*(n=3) and *Ntv-a;Trem2^-/-^* (n=3) *Nf1* tumors. **(H)** UMAP dimensionality reduction and subclustering of the myeloid populations from the entire dataset. **(I)** UMAP subclustering of the microglial population from the entire dataset. **(J)** Dot plot highlighting the top 5 marker genes expressed by each of the 5 microglial subclusters. **(K and L)** UMAP subclustering of the monocyte-derived macrophage (MDM) and monocyte populations respectively from the entire dataset. **(M)** Bar plot showing the proportion of the 5 microglial subclusters between *Ntv-a;Trem2^+/+^* (n=3) and *Ntv-a;Trem2^-/-^*(n=3) *Nf1* tumors. **(N)** Dot plot indicating the percent expression and average expression of *Trem2* across the 5 microglial subclusters outlined in (I & M). **(O-S)** Box plots comparing the various subclusters of microglia, MDMs and monocytes between *Ntv-a;Trem2^+/+^*(n=3) and *Ntv-a;Trem2^-/-^* (n=3) *Nf1* tumors as a percentage of the respective parent populations. Wilcoxon rank-sum test *p<0.05, **p<0.01.

Since our analysis of scRNA-seq data across murine GBM models with different driver mutations highlighted *Trem2* expression primarily in the microglial and MDM subclusters (Fig. S1B), we next investigated changes in these populations in *Trem2* WT and deficient *Nf1* mGBM. We first subclustered the total myeloid subpopulation (Fig. 5H), comprising 23,517 cells and representing the dominant myeloid subsets in GBM such as microglia, MDMs, and monocytes, along with less abundant subsets such as neutrophils, DCs, and plasmacytoid DCs (pDCs). From this myeloid population, we then isolated microglial, MDM, and monocyte subclusters based on our previously published marker genes for each cell type (Fig. 5I-L and Fig. S11A&D). Across the microglial subclusters, we observed the highest expression of *Trem2* in homeostatic microglia, followed closely by disease-associated and pro-inflammatory microglia (Fig. 5N). *Trem2* was also highly expressed in the disease-associated, homeostatic, and pro-inflammatory MDMs (Fig. S11B). Homeostatic microglia, expressing markers such as *P2ry12*, exhibited a significant increase in the *Trem2*-deficient tumors (Fig. 5M&O). Perhaps the most significant finding was the decrease in disease-associated and pro-inflammatory microglia in *Trem2*-deficient tumors (Fig. 5M, P & Q). Moreover, this trend of an increase in the homeostatic cell state along with a decrease in the pro-inflammatory state was conserved across the MDM and monocyte populations (Fig. 5R&S and Fig. S11C, E). Overall, these changes suggest that *Trem2* supports disease-associated and pro-inflammatory myeloid states, while *Trem2* deficiency favors homeostatic states. These changes are associated with survival extension in *Trem2*-deficient *Nf1* mGBM-bearing mice (Fig. 1B) and, together with the tumor secretome findings (Fig. 4E), suggest a role for TREM2 in shaping myeloid states in Ras-driven GBM.

Beyond the tumor and myeloid compartments, we observed modest changes in the astrocyte and endothelial populations in *Trem2*-deficient *Nf1* mGBM (Fig. 5B-D). Using a previously published framework to subcluster the astrocytes^80^, we identified the mature astrocyte signature as the predominant program (Fig. S12A-D). IHC staining showed an increase in GFAP-positive area within *Trem2*-deficient tumors (Fig. S13C&D). Endothelial cell subclustering showed a reduction in Angpt2^+^, proliferating, and interferon-associated clusters alongside expansion of a Cxcl12^+^ cluster (Fig. S12E-H). However, CD31^+^ vascular coverage and mean vessel size were comparable between genotypes (Fig. S13E&F). Together, these findings suggest that *Trem2* deficiency is associated with transcriptional changes in both astrocytes and endothelial cells, although without major detectable changes in the measured vascular parameters.

### Early deletion of *Trem2* in brain-resident myeloid cells prolongs survival in Ras oGBM

Because the effects of TREM2 signaling are highly context-dependent, we next aimed to determine whether the tumor-supportive role of *Trem2* in Ras oGBM depends on the myeloid compartment in which it is expressed and the timing of its activity during tumor progression. To distinguish the contributions of brain-resident and infiltrating myeloid populations, we generated bone marrow chimeras with lead head shielding to replace circulating myeloid cells while protecting those resident to the CNS (Fig. 6A). We used *Trem2^-/-^* and CD45.1 WT (PepBoy) mice to create chimeric mice and distinguish donor versus recipient cells. Our results showed that *Trem2^-/-^* recipients reconstituted with PepBoy bone marrow had smaller tumor burden compared to PepBoy recipients of *Trem2^-/-^*bone marrow (Fig. 6B). These results indicate that brain-resident microglia-specific loss of *Trem2* is responsible for slowing Ras-driven oGBM progression.

**Figure 6:**
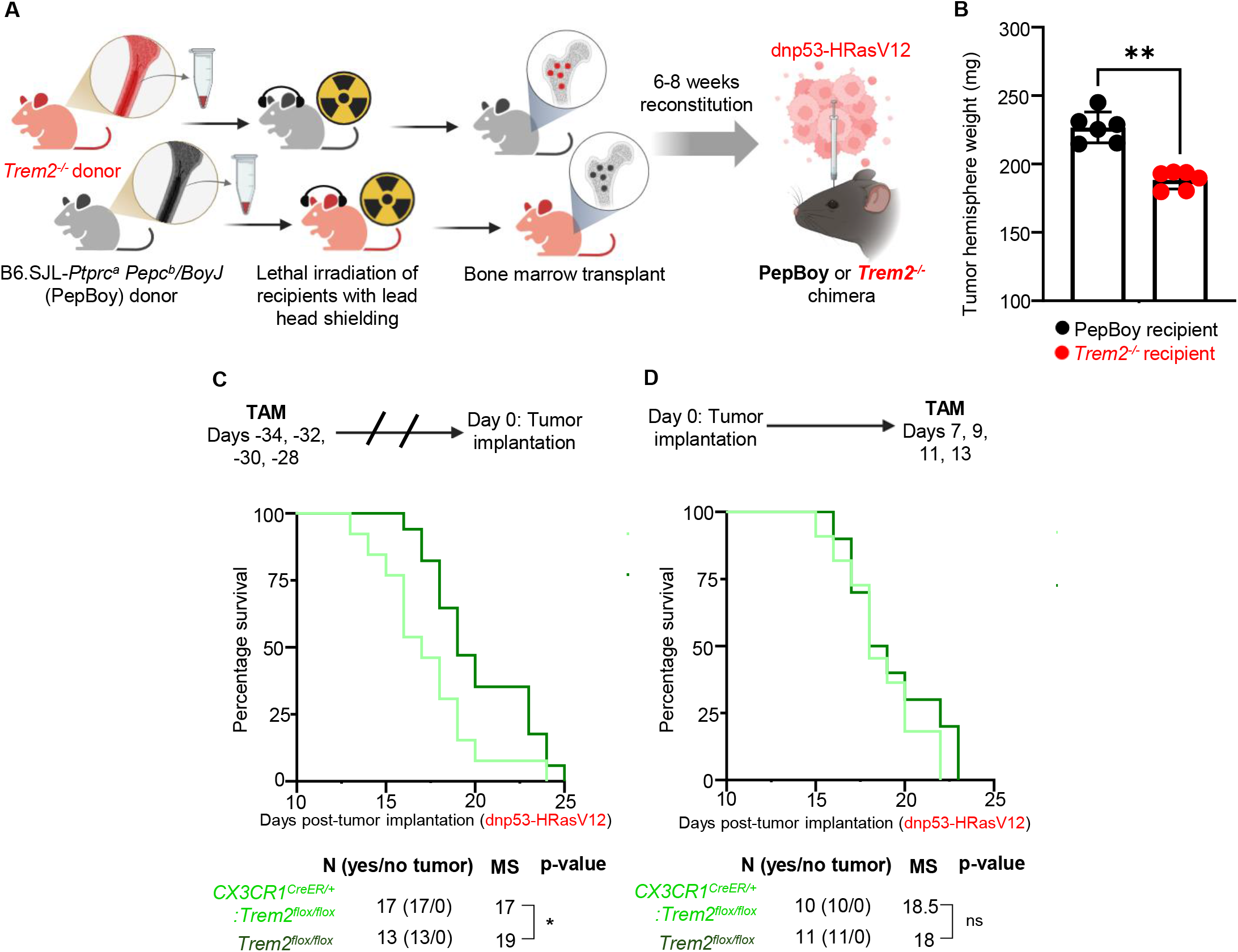
Early-stage, microglial *Trem2* is sufficient for the phenotype in Ras-driven oGBM. **(A)** Schematic depicting generation of bone marrow chimeric mice. Created in Biorender. **(B)** Tumor-bearing hemisphere weight of PepBoy (n=6) and *Trem2^-/-^*(n=6) chimeric mice with Ras-driven oGBM. Two-tailed, unpaired t-test. **(C)** Tamoxifen administration timeline and Kaplan-Meier survival curve for early *Trem2* conditional knockout (*CX3CR1^CreER/+^Trem2^flox/flox^*, n=17) versus *Trem2^flox/flox^* control (n=13) with Ras-driven oGBM. Gehan-Breslow-Wilcoxon test. **(D)** Tamoxifen administration timeline and Kaplan-Meier survival curve for late-stage *Trem2* conditional knockout (*CX3CR1^CreER/+^Trem2^flox/flox^*, n=10) versus *Trem2^flox/flox^* control (n=11) with Ras-driven oGBM. Gehan-Breslow-Wilcoxon test. **(E)** TAM=tamoxifen, MS=median survival; *p<0.05, **p<0.01.

To more directly examine the contribution of microglial TREM2 using a genetic approach and determine whether the effects of *Trem2* loss depended on the timing of deletion, we crossed *Trem2^flox/flox^* mice with *CX3CR1^CreER/+^* mice to generate an inducible conditional knockout model. We confirmed successful deletion of *Trem2* after tamoxifen administration (Fig. S14A&B). In the early deletion paradigm, tamoxifen was injected a month prior to tumor implantation, allowing recombined circulating monocytes to turn over while retaining recombination in long-lived microglia.^81^ Deletion of *Trem2* before implantation replicated the survival benefit observed following germline *Trem2* loss (Fig. 6C). In contrast, tamoxifen administration after tumors were established did not change the median survival (Fig. 6D). Together, these findings support an early contribution of TREM2 in microglia to Ras-driven oGBM progression and indicate that deletion before tumor implantation, but not after tumor establishment, confers a survival benefit.

## Discussion

In this study, we show that genetic driver mutations influence the effects of *Trem2* deficiency on tumor progression and the GBM TME. Across *de novo* and orthotopic murine GBM models, we find that *Trem2* deficiency improves survival and tumor burden in Ras oGBM and *Nf1* mGBM, whereas it increased tumor burden without affecting survival in PDGFB-driven GBM. These different outcomes were accompanied by distinct changes in the myeloid compartment and soluble factor profiles. *Trem2* deficiency reduced microglial abundance in *Nf1* mGBM and Ras oGBM, but decreased infiltrating BMDM, monocytes, and neutrophils in PDGFB-driven tumors. These differences align with the genetic driver-associated myeloid composition we identified in our previous single-cell analyses^16,17^ and suggest that the consequences of *Trem2* loss depend, in part, on the cellular context in which it functions. The soluble-factor changes associated with *Trem2* deficiency also differed between models. Ras-driven tumors showed higher levels of inflammatory and lymphocyte-recruiting mediators, including CXCL9 and CXCL10, but these changes were not accompanied by increased T-cell abundance at the tumor endpoint. In contrast, the PDGFB oGBM TME displayed increases in immunosuppressive factors, including IL-6, CXCL1, and IGFBP-1. Together, these findings indicate that *Trem2* deficiency remodels the TME in a genetic driver-dependent manner, increasing inflammatory mediators in Ras oGBM while producing a mixed profile of immune-regulatory and tumor-associated factors in PDGFB oGBM.

We found that pBMDM migration and microglial process extension/retraction depended on TREM2 in response to Ras-derived, but not PDGFB-derived TCM. Ras-derived TCM was enriched in proteins associated with disease-associated myeloid states and matrix remodeling, whereas PDGFB-derived TCM was enriched in neural extracellular matrix proteins. Among the Ras-enriched factors, SPP1 and fibronectin (FN1) may be particularly relevant given their known roles in cytoskeletal remodeling, integrin-mediated adhesion, and myeloid cell migration.^82,83^ Conversely, the neural extracellular matrix components in PDGFB-derived TCM may establish different factors for myeloid cell motility or process extension. Although these secretome differences identified candidates that may contribute to TREM2-dependent myeloid responses, their direct interaction with TREM2 and functional roles remain to be tested. The increase in tumor burden without a survival difference in PDGFB-driven GBM-bearing mice may reflect a growth effect that is insufficient to change overall survival. Similar findings have been reported in GL261 and CT2A models, where *Trem2* deficiency increased tumor burden or accelerated growth without changing survival.^36,37^ Another study has reported tumor-promoting effects of TREM2, including sex-dependent effects on GL261 growth and angiogenesis, further emphasizing the influence of experimental context.^35^

A major finding of our single-cell analysis of *Nf1* mGBM was the shift in myeloid transcriptional states following *Trem2* loss. *Trem2-*deficient *Nf1* mGBM showed reduced proportions of disease-associated and pro-inflammatory microglia and increased homeostatic microglia. We observed a similar shift away from pro-inflammatory states and toward homeostatic states in MDMs and monocytes, indicating that *Trem2* deficiency affects multiple myeloid populations. These findings parallel the secretome analyses as well as our previous observations in diffuse midline glioma (DMG), where a shift from disease-associated to homeostatic microglia was associated with improved survival.^84^ Interestingly, this contrasts with neurodegeneration, where TREM2-dependent transitions from homeostatic to disease-associated microglia are normally considered protective.^26^ In GBM, however, these same disease-associated cellular programs may instead drive immunosuppression, while homeostatic microglia are able to resist tumor progression through acquisition of a unique transcriptional program capable of producing pro-inflammatory factors.

Together, these results suggest that TREM2 supports disease-associated and inflammatory myeloid programs in *Nf1* mGBM and that the shift toward homeostatic states may contribute to the survival benefit observed in *Trem2*-deficient GBM-bearing mice. Whether *Trem2*deficiency similarly alters disease-associated and homeostatic myeloid states in PDGFB-driven GBM remains to be determined. However, distinct proportions of microglia and MDM in PDGFB tumors and TREM2-dependent increases in immunosuppressive factors in the PDGFB oGBM TME suggest that TREM2 may support different functional reprogramming based on genetic drivers.

Our bone marrow chimera experiments support a predominant role for TREM2 from brain-resident microglia and potentially skull marrow-derived myeloid cells, consistent with a microglial contribution to the phenotype in Ras oGBM. We further supported these findings using a conditional genetic approach. Tamoxifen administration one month before tumor implantation allowed recombined circulating monocytes to turn over while retaining *Trem2* deletion in brain-resident myeloid cells, including microglia.^81^ This early deletion prolonged survival in mice bearing Ras oGBM. In contrast, tamoxifen administration after tumor establishment, which targeted *Trem2* deletion in both microglia and infiltrating monocytes and MDMs, did not confer a detectable survival benefit. Together, these results support an early contribution of brain-resident myeloid TREM2 to Ras-driven GBM progression. The stronger effect of *Trem2* loss in *Nf1* mGBM and Ras oGBM may also reflect their greater microglial content relative to PDGFB-driven tumors, consistent with the enrichment of microglia reported in human MES GBM.^48^ In line with this, we have recently shown that H3.1K27 DMG are enriched in BMDMs, whereas H3.3K27M DMG are enriched in microglia, and that blocking BMDM recruitment or depleting BMDMs only impacted the survival of H3.1K27M DMG, demonstrating differential myeloid dependence.^45^

TREM2 is a promising therapeutic target in many neurological disorders and is currently in clinical trials for Alzheimer’s disease.^85,86^ However, the context-dependent nature of TREM2 suggests that therapeutic outcomes could be influenced by disease stage, cellular identity, and the brain microenvironment. Our work identifies genetic driver mutations as an additional determinant of TREM2 function in GBM, which carries implications for identifying patients most likely to benefit from TREM2-directed therapy.

Together, our findings show that TREM2 targeting is not uniformly beneficial across GBM, suggesting that general TREM2-activating or inhibiting strategies may not be efficacious for all patients. Instead, these results highlight tumor genotype, myeloid composition, and timing as key considerations for TREM2-directed therapies.

## Acknowledgments

We thank all members of the Wu and Hambardzumyan labs for scientific discussion of the results. We also thank the UTHealth Houston Cancer Genomics Center, MD Anderson ORION (Oncology Research and Immuno-mONitoring) Core, the UTHealth Houston Clinical and Translational Proteomics Center, and the UTHealth Center for Advanced Microscopy (a Nikon Center of Excellence) for their technical assistance. We thank the Flow Cytometry Core and the Human Immune Monitoring Core at the Icahn School of Medicine at Mount Sinai for providing resources and assistance for spectral flow cytometry and single-cell RNA sequencing. We additionally thank the Penn Brain Tumor Center at the Perelman School of Medicine, University of Pennsylvania, for providing resources for confocal microscopy and immunohistochemistry. We thank Dr. Q Richard Lu and Dr. Marco Colonna for providing cell lines and *Trem2^-/-^*mice. We also thank Dr. Rodney Ritzel, Dr. Chris Janssen, and Ruby Janssen for assistance with irradiation experiments, and Dr. Andrea Becerra Calixto and Dr. Louise McCullough for their support in efferocytosis experiments.

## Supplementary material

Supplementary material is available online.

## Data availability

Data will be made available upon request.

---

**Figure S1: T*rem2* is expressed in GBM-associated microglia and macrophages independent of genetic driver mutation. (A)** Schematic depicting *Nf1* (n=3, red), *PDGFB* (n=4, blue), and *EGFRvIII* (n=3, green) mGBM analyzed from the Soni, *et al.* single cell atlas. Created in Biorender. **(B)** Dot plot showing percent expression and average expression of *Trem2* across myeloid populations in wildtype *Nf1, PDGFB,* and *EGFRvIII* tumors. **(C)** Bar plots showing percent *Trem2^+^* cells in total myeloid, microglia, and MDM populations across the three tumor models. **(D)** Bar plots showing expression of a *Trem2* pathway module score in total myeloid, microglia, and MDM populations across the three tumor models.

**Figure S2: Validation of TREM2 expression in *Trem2-*deficient and wild-type *Ntv-a* mice. (A)** Plot showing *Trem2* transcript levels in scRNAseq data compared between n=3 *Ntv-a;Trem2^+/+^* and n=3 *Ntv-a;Trem2^-/-^ Nf1* tumors. Wilcoxon rank-sum test ****p<0.0001 **(B)** IF co-staining for TREM2 (red) and IBA1 (green) in *Ntv-a;Trem2^+/+^* and *Ntv-a;Trem2^-/-^* brain sections. DAPI (blue) was used as a nuclear counterstain. Scale bar = 10 μm.

**Figure S3: Immune profiling of *Nf1* and *PDGFB*-driven tumors from *Trem2*-deficient and wild-type mice. (A and B)** Analysis of phenotypic markers MHC-II and PDL1 in microglia and MDMs from *Nf1* tumors compared between *Ntv-a;Trem2^+/+^* (n=10) and *Ntv-a;Trem2^-/-^* (n=9) mice as a percentage of live cells (left) and as a heatmap of mean fluorescent intensity (MFI) (right). Two-tailed, unpaired t-test. **(C and D)** Analysis of phenotypic markers MHC-II and PDL1 in microglia and MDMs from *PDGFB*-driven tumors compared between *Ntv-a;Trem2^+/+^*(n=5) and *Ntv-a;Trem2^-/-^* (n=5) mice as a percentage of live cells (left) and as a heatmap of mean fluorescent intensity (MFI) (right). Two-tailed, unpaired t-test. **(E)** Plots of various lymphoid populations in *Nf1* tumors compared between *Ntv-a;Trem2^+/+^* (n=9) and *Ntv-a;Trem2^-/-^* (n=9) mice as a percentage of live cells. Two-tailed, unpaired t-test. **(F)** Plots of various lymphoid populations in *PDGFB*-driven tumors compared between *Ntv-a;Trem2^+/+^* (n=5) and *Ntv-a;Trem2^-/-^* (n=5) mice as a percentage of live cells. Two-tailed, unpaired t-test. **(G&H)** Analysis of phenotypic markers PD1 and TIM3 in CD4^+^ and CD8^+^ T cells from *Nf1* tumors compared between *Ntv-a;Trem2^+/+^* (n=9) and *Ntv-a;Trem2^-/-^* (n=9) mice as a percentage of live cells (left) and as a heatmap of mean fluorescent intensity (MFI) (right). Two-tailed, unpaired t-test. **(I&J)** Analysis of phenotypic markers PD1 and TIM3 in CD4^+^ and CD8^+^ T cells from *PDGFB*-driven tumors compared between *Ntv-a;Trem2^+/+^* (n=5) and *Ntv-a;Trem2^-/-^* (n=5) mice as a percentage of live cells (left) and as a heatmap of mean fluorescent intensity (MFI) (right). Two-tailed, unpaired t-test. *p<0.05.

**Figure S4: Flow cytometry immune profiling of wild-type and *Trem2^-/-^* mice with oGBM. (A)** Proportions of myeloid populations in wild-type and *Trem2^-/-^* Ras-driven oGBM 14 days-post tumor implantation. Two-tailed, unpaired t-test. **(B)** Proportions of lymphoid populations in wild-type and *Trem2^-/-^* Ras-driven oGBM 14 days-post tumor implantation. Two-tailed, unpaired t-test. **(C)** Proportion of myeloid populations in wild-type and *Trem2^-/-^*PDGFB-driven oGBM 21 days-post tumor implantation. Two-tailed, unpaired t-test. **(D)** Proportion of lymphoid populations in wild-type and *Trem2^-/-^* PDGFB-driven oGBM 21 days-post tumor implantation. Two-tailed, unpaired t-test.*p<0.05.

**Figure S5: Tre*m1* expression is reduced in *Trem2*-deficient mice but is not sufficient to drive survival and immune remodeling in *Nf1* and *PDGFB* mGBM. (A)** Dot plot of *Trem1* expression from the scRNAseq dataset of *Ntv-a;Trem2^+/+^* (n=3) and *Ntv-a;Trem2^-/-^*(n=3) *Nf1* mGBM. **(B)** Violin plot indicating the expression of *Trem1* across all cell populations in the scRNAseq dataset. **(C)** Schematic of *de novo Nf1*-driven (red) and *PDGFB*-driven (blue) mGBM generated in *Ntv-a;Trem1^+/+^* and *Ntv-a;Trem1^-/-^*mice. Created in Biorender. **(D)** Kaplan-Meier curve comparing survival of *Ntv-a;Trem1^+/+^*(n=9) and *Ntv-a;Trem1^-/-^* (n=10) mice with *PDGFB* tumors; Gehan-Breslow-Wilcoxon test. **(E)** Kaplan-Meier curve comparing survival of *Ntv-a;Trem1^+/+^* (n=35) and *Ntv-a;Trem1^-/-^* (n=27) mice with *Nf1* tumors; Gehan-Breslow-Wilcoxon test. **(F and G)** Plots showing flow cytometry analysis of various myeloid and lymphoid immune populations compared between *Ntv-a;Trem1^+/+^* (n=6) and *Ntv-a;Trem1^-/-^* (n=6) *PDGFB* mGBM. Two-tailed, unpaired t-test. **(H and I)** Plots showing flow cytometry analysis of various myeloid and lymphoid immune populations compared between *Ntv-a;Trem1^+/+^*(n=15) and *Ntv-a;Trem1^-/-^* (n=14) *Nf1* tumors. Two-tailed, unpaired t-test. *p<0.05.

**Figure S6: Cytokine profiling array of *Trem2^-/-^*and wild-type PDGFB oGBM. (A)** Waterfall plot displaying fold changes for cytokines and chemokines in wild-type and *Trem2^-/-^* PDGFB oGBM tumor lysates.

**Figure S7: Microglial phagocytosis of GSCs is impaired by *Trem2* deficiency regardless of tumor genotype. (A)** Proportion of *CX3CR1^GFP/+^* (n=5) and *CX3CR1^GFP/+^Trem2^-/-^*(n=5) microglia with tdTomato^+^ inclusions after co-culture with apoptotic dnp53-HRasV12-tdTomato cells. Two-way ANOVA. **(B)** Proportion of *CX3CR1^GFP/+^* (n=5) and *CX3CR1^GFP/+^Trem2^-/-^* (n=5) microglia with tdTomato^+^ inclusions after co-culture with apoptotic dnp53-PDGFB-tdTomato cells. Two-way ANOVA. **p<0.01, ***p<0.001.

**Figure S8: Bulk RNA sequencing of CD11b^+^ myeloid cells sorted from wild-type and *Trem2^-/-^* Ras and PDGFB oGBM.** Volcano plots showing differentially expressed genes between **(A)** WT and *Trem2^-/-^* Ras-driven oGBM, **(B)** WT and *Trem2^-/-^* PDGFB-driven oGBM, **(C)** PDGFB and Ras-driven oGBM from WT mice, and **(D)** PDGFB and Ras-driven oGBM from *Trem2^-/-^* mice. Significance is defined by adjusted p-value <0.05 and |log2 fold change| <1.

**Figure S9: Single-cell RNA sequencing analysis of *Trem2*-deficient and wild-type *Nf1* mGBM. (A)** Dot plot depicting the top marker genes defining each cell population within the entire dataset. **(B)** UMAP dimensionality reduction of cycling and non-cycling cells within the entire dataset. **(C)** Bar plot showing the proportion of cells in G2M, S and G1 phases of the cell cycle compared between n=3 *Ntv-a;Trem2^+/+^* and n=3 *Ntv-a;Trem2^-/-^ Nf1* tumors. **(D)** Box plots showing the various subpopulations respectively in *Ntv-a;Trem2^+/+^* (n=3) and *Ntv-a;Trem2^-/-^* (n=3) *Nf1* tumors as a percentage of all cells.

**Figure S10: Single cell sequencing analysis of tumor cell states in *Trem2*-deficient and wild-type *Nf1* mGBM. (A)** UMAP dimensionality reduction of *UC3-R-RCAS* expression within the tumor subpopulation of the scRNAseq dataset. **(B)** UMAP dimensionality reduction of cycling and non-cycling cells within the tumor subpopulation of the scRNAseq dataset. **(C)** UMAP dimensionality reduction of the module scores of the four Neftel GBM cell states projected onto the tumor subpopulation of the scRNAseq dataset. **(D)** Box plots of the mean module scores of the four Neftel GBM cell states compared between the tumor compartment of n=3 *Ntv-a;Trem2^+/+^* and n=3 *Ntv-a;Trem2^-/-^* samples. Wilcoxon rank-sum test. **(E)** Dot plot depicting differentially expressed genes within the tumor compartment of n=3 *Ntv-a;Trem2^+/+^* and n=3 *Ntv-a;Trem2^-/-^* samples.

**Figure S11: Single cell sequencing analysis of the MDM and monocyte subpopulations in *Trem2*-deficient and wild-type *Nf1* mGBM. (A)** Dot plot highlighting the top 5 marker genes expressed by each of the 5 MDM subclusters. **(B)** Dot plot indicating the percent expression and average expression of *Trem2* across the 5 MDM subclusters. **(C)** Box plots comparing the various subclusters of MDMs between *Ntv-a;Trem2^+/+^* (n=3) and *Ntv-a;Trem2^-/-^* (n=3) *Nf1* tumors as a percentage of the total MDM population. Wilcoxon rank-sum test. **(D)** Dot plot highlighting the top 5 marker genes expressed by each of the 3 monocyte subclusters. **(E)** Box plots comparing disease-associated and pro-inflammatory monocyte subclusters between *Ntv-a;Trem2^+/+^* (n=3) and *Ntv-a;Trem2^-/-^* (n=3) *Nf1* tumors as a percentage of the total monocyte population. Wilcoxon rank-sum test.

**Figure S12: Single cell sequencing analysis of the astrocyte and endothelial cell subpopulations in *Trem2*-deficient and wild-type *Nf1* mGBM. (A)** UMAP dimensionality reduction of the entire single cell dataset with the astrocyte subpopulation circled. (B) Astrocyte subclustering and UMAP dimensionality reduction of the inflammatory, alternative activation, progenitor-like and mature astrocyte modules from Heiland, *et al.*^75^ projected onto the dataset. (C) Two-dimensional quadrant scatter plot of the four astrocyte states defined in (B) compared between *Ntv-a;Trem2^+/+^* (n=3) and *Ntv-a;Trem2^-/-^*(n=3) *Nf1* tumors. (D) Bar plot showing the proportion of the four astrocyte states compared between *Ntv-a;Trem2^+/+^* (n=3) and *Ntv-a;Trem2^-/-^* (n=3) *Nf1* tumors. (E) UMAP dimensionality reduction and subclustering of the endothelial cell population from the dataset. (F) Dot plot indicating the top marker genes associated with each endothelial cell subcluster. (G) Bar plot showing the proportion of each endothelial cell subcluster from the UMAP in (E) compared between *Ntv-a;Trem2^+/+^* (n=3) and *Ntv-a;Trem2^-/-^* (n=3) *Nf1* tumors. (H) Box plots comparing the seven endothelial cell subclusters between *Ntv-a;Trem2^+/+^* (n=3) and *Ntv-a;Trem2^-/-^* (n=3) *Nf1* tumors. Wilcoxon rank-sum test.

**Figure S13: Immunohistochemical staining for various markers in *Trem2*-deficient and wild-type *Nf1* mGBM. (A)** Representative images and **(B)** bar plot indicating the quantification of pHH3^+^ cells per square millimetre of tumor in *Ntv-a;Trem2^+/+^* (n=7) and *Ntv-a;Trem2^-/-^* (n=6) *Nf1* tumors. Values were compared by two-tailed, unpaired t-test. **(C)** Representative images and **(D)** bar plot indicating quantification of GFAP^+^ area within the tumors of *Ntv-a;Trem2^+/+^*(n=6) and *Ntv-a;Trem2^-/-^* (n=6) mice. Each dot represents the average of values from 5 representative images per sample. Values were compared by two-tailed, unpaired t-test.**p<0.01. Scale bar = 50 μm. **(E)** Representative images and **(F)** bar plots indicating quantification of CD31^+^ area (left) and average vessel size (right) within the tumors of *Ntv-a;Trem2^+/+^* (n=6) and *Ntv-a;Trem2^-/-^* (n=6) mice. Each dot represents the average of values from 5 representative images per sample. Values were compared by two-tailed, unpaired t-test.

**Figure S14: Validation of the conditional *Trem2* knockout mouse model. (A)** Graphic showing isolation of CD11b^+^ cells from naïve *CX3CR1^CreER/+^Trem2^flox/flox^* conditional knockout and control *Trem2^flox/flox^* mice after tamoxifen administration. Created in Biorender. **(B)** rt-qPCR gene expression of *Trem2* in conditional knockout normalized to *Trem2^flox/flox^* control mice (n=4 per group). Unpaired, two-tailed t-test, **p<0.01.

