## Supplementary Figures for "Genetic drivers shape TREM2-dependent myeloid responses and glioblastoma progression"

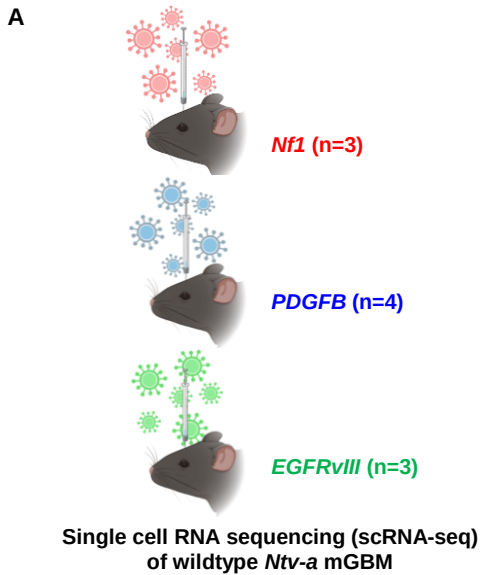

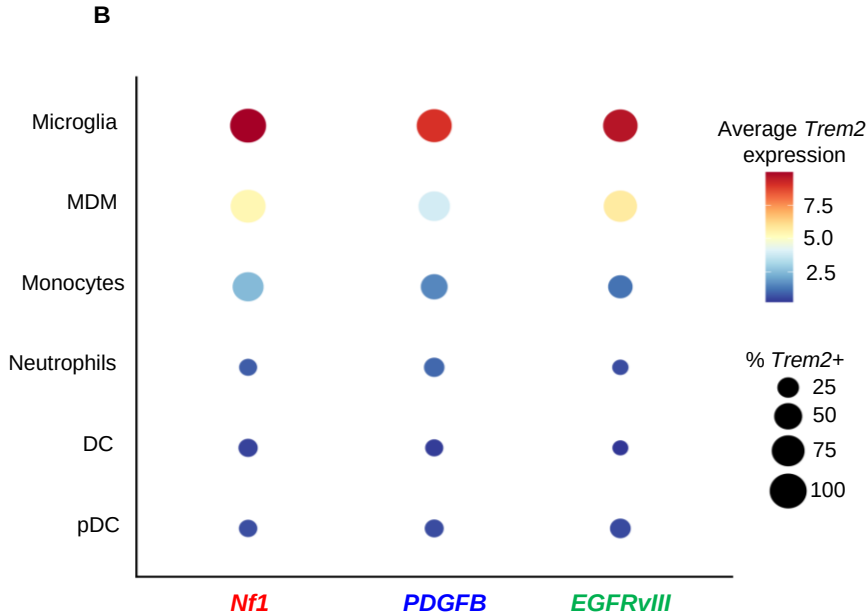

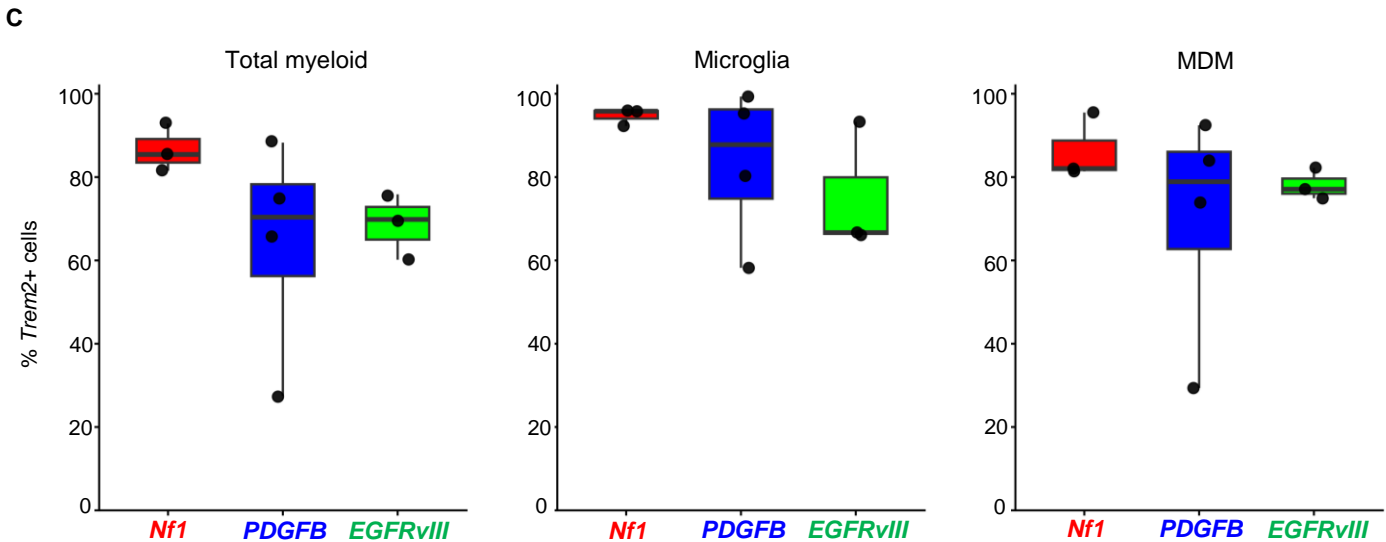

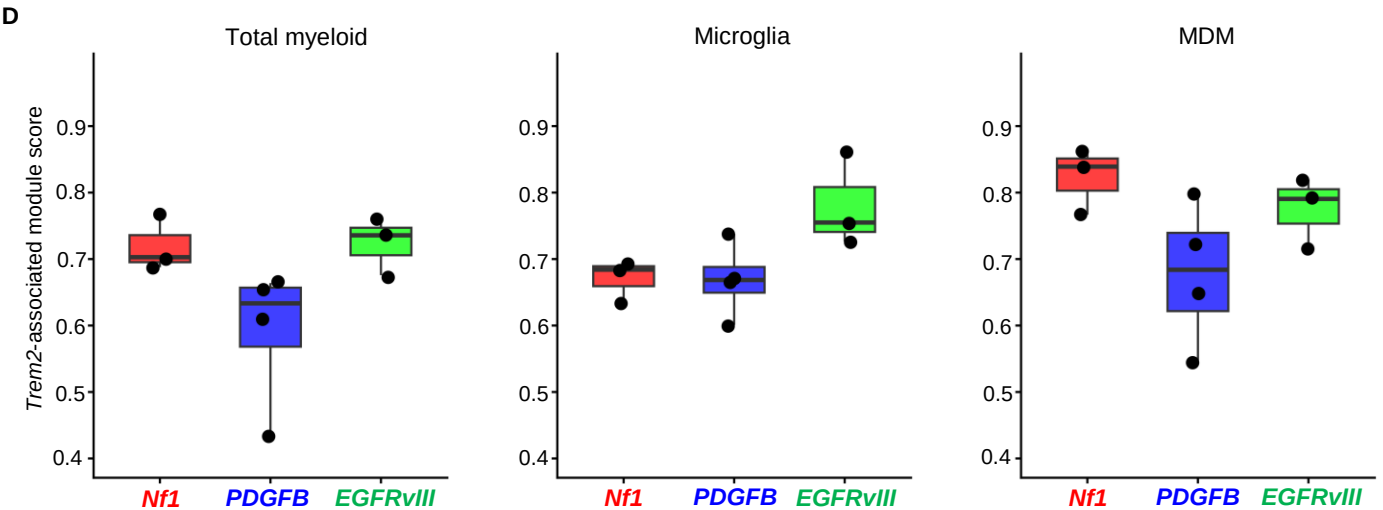

A

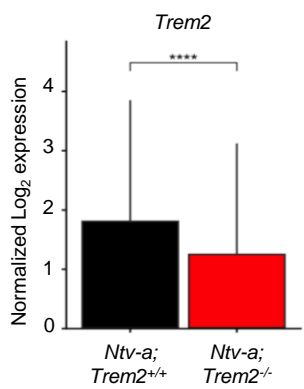

B

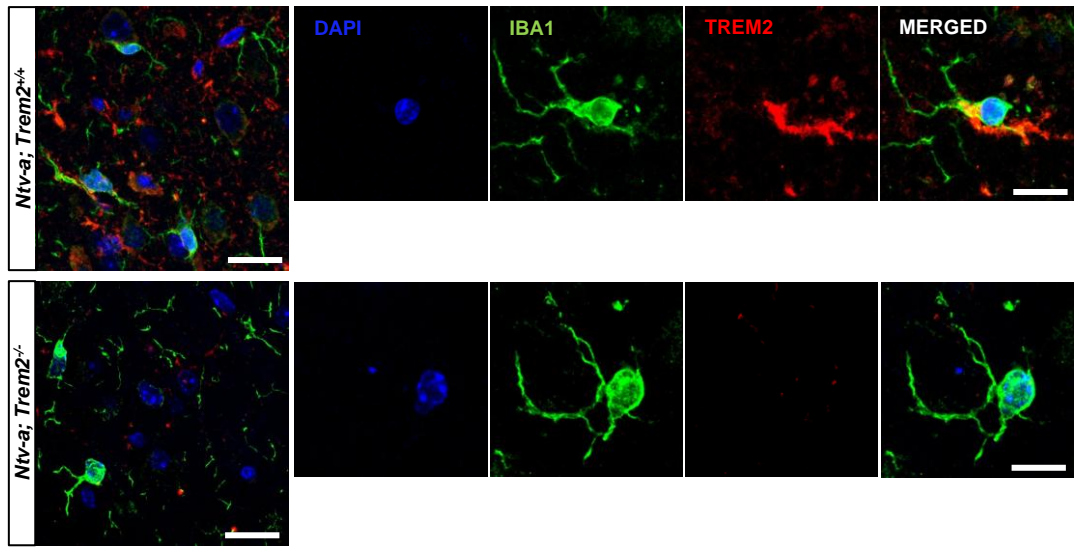

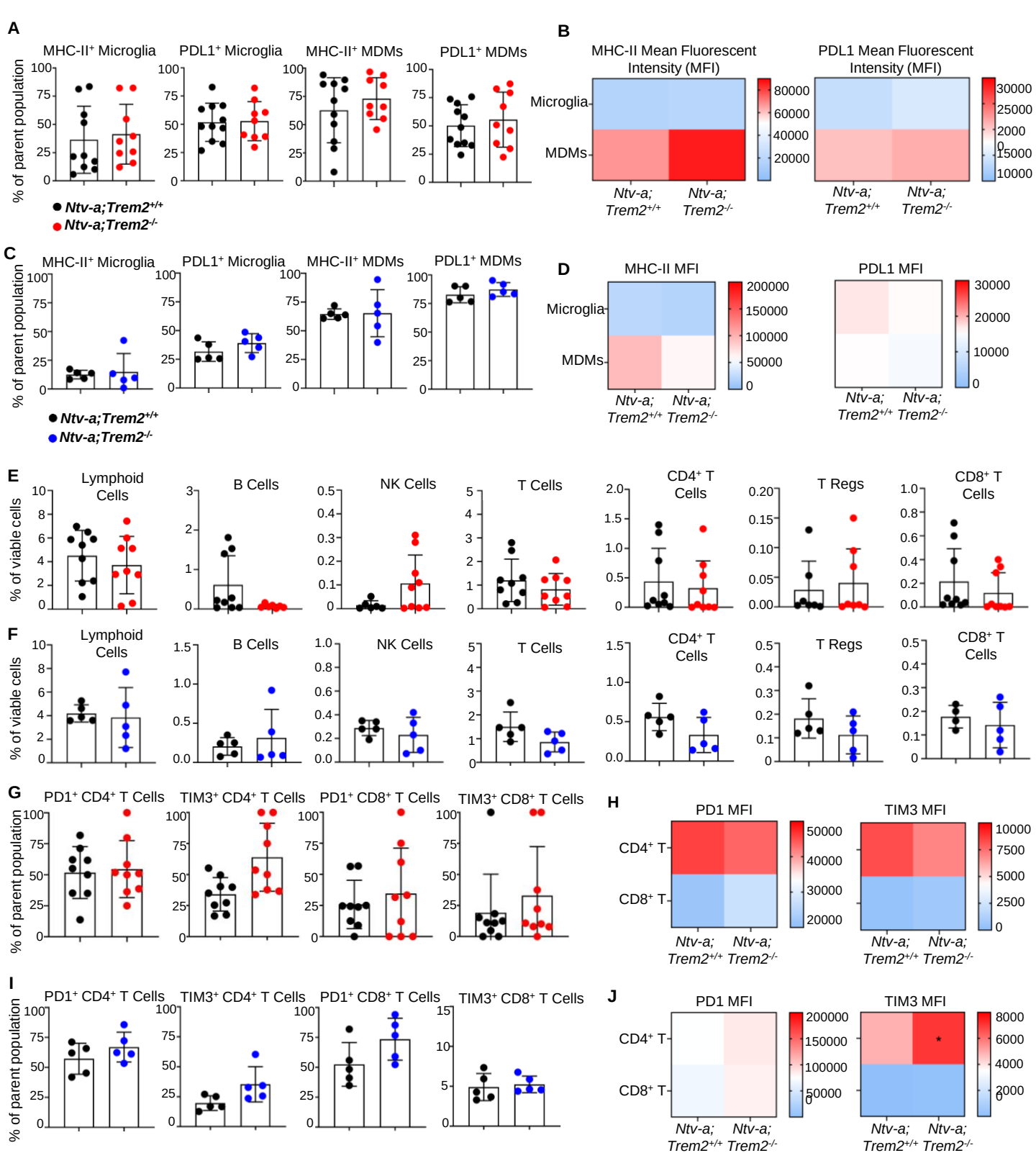

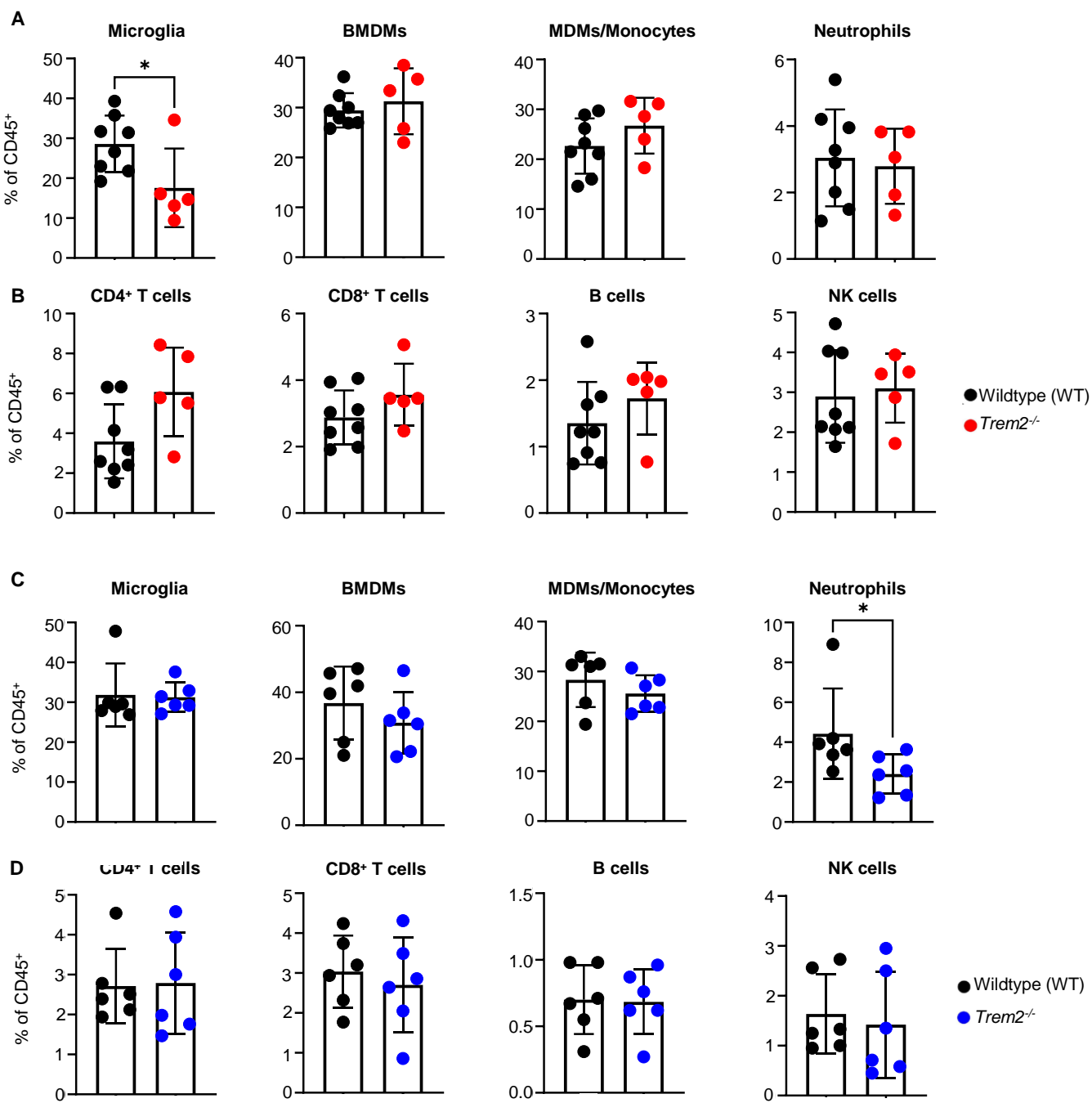

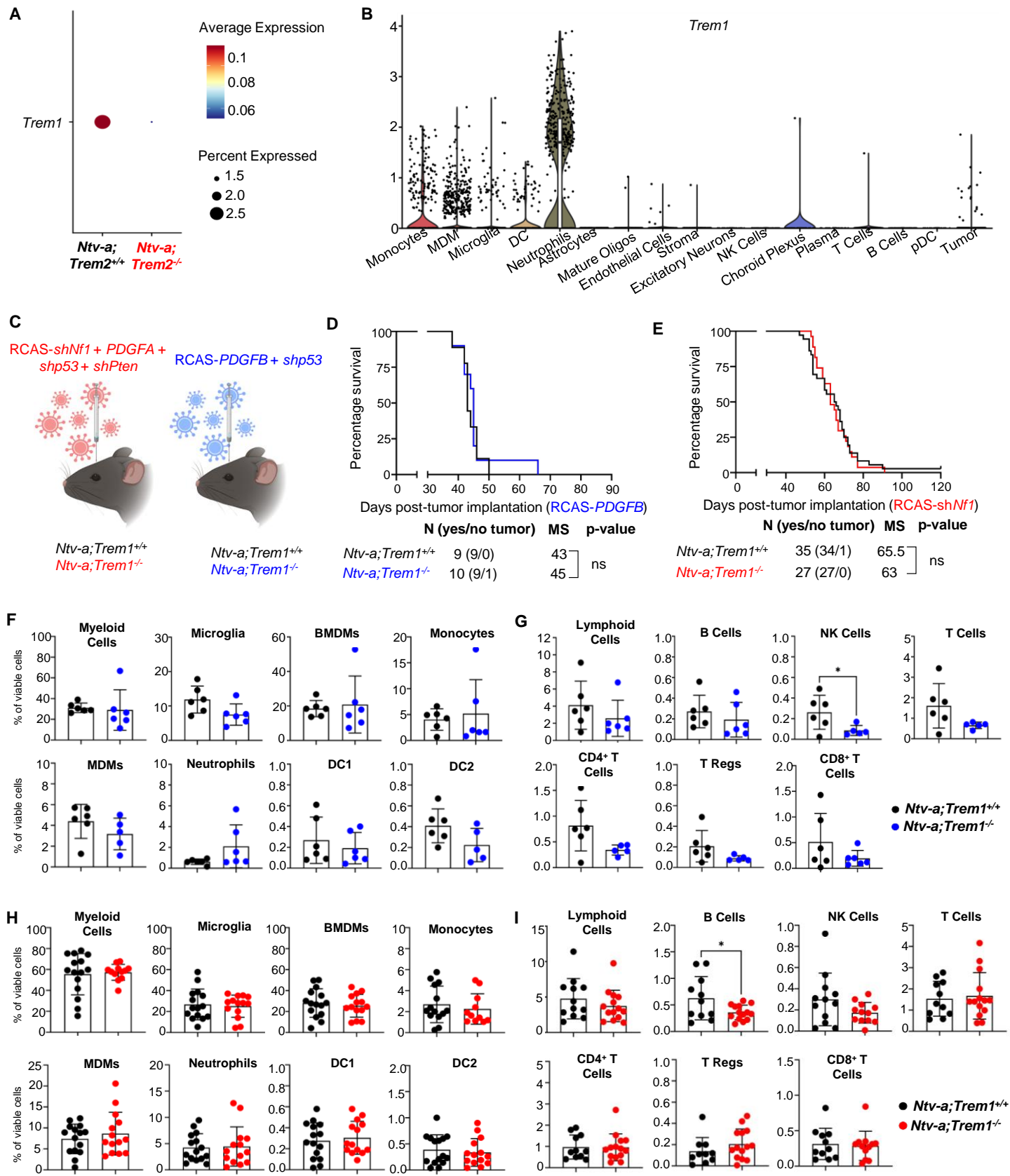

A

Up in Trem2<sup>-/-</sup> Down in Trem2<sup>-/-</sup>

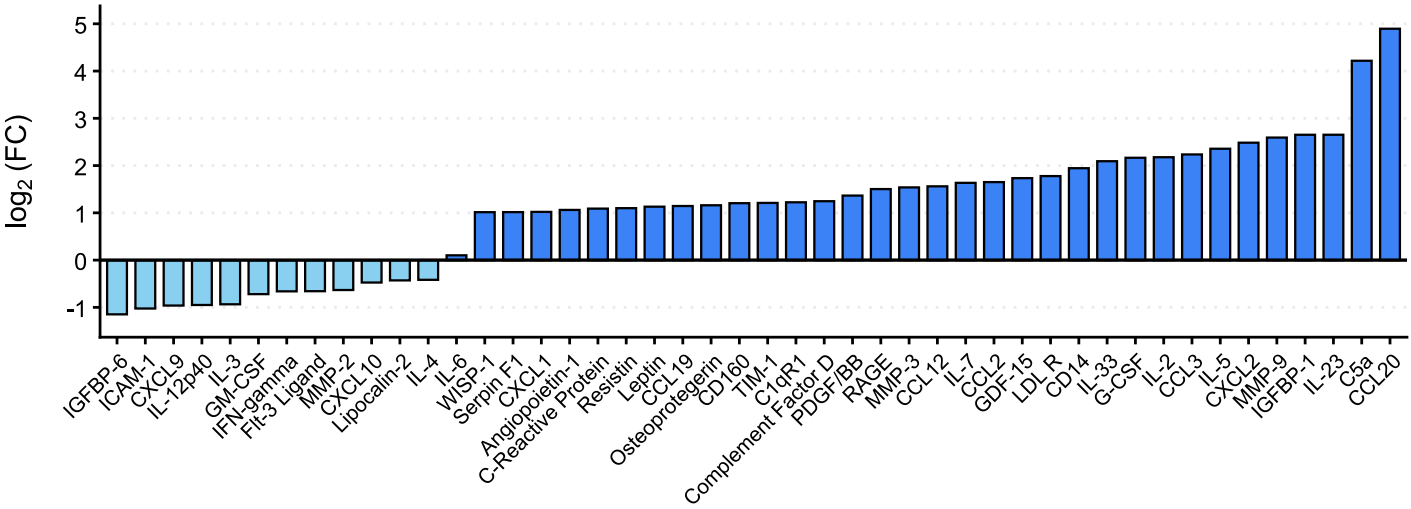

A

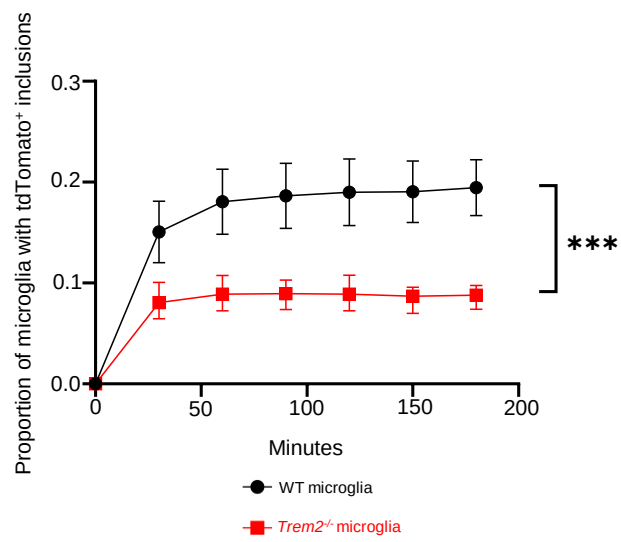

B

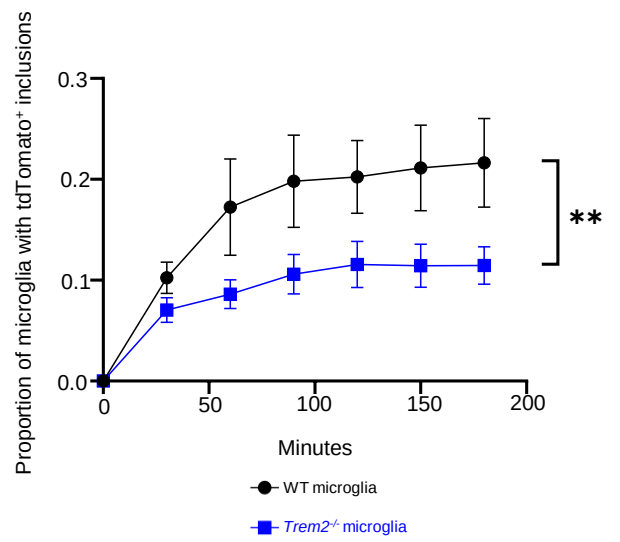

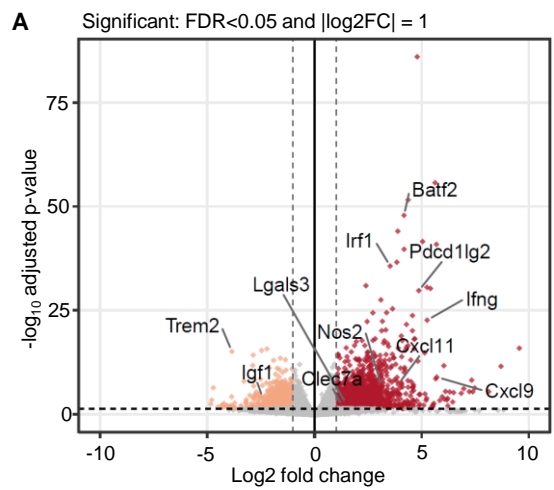

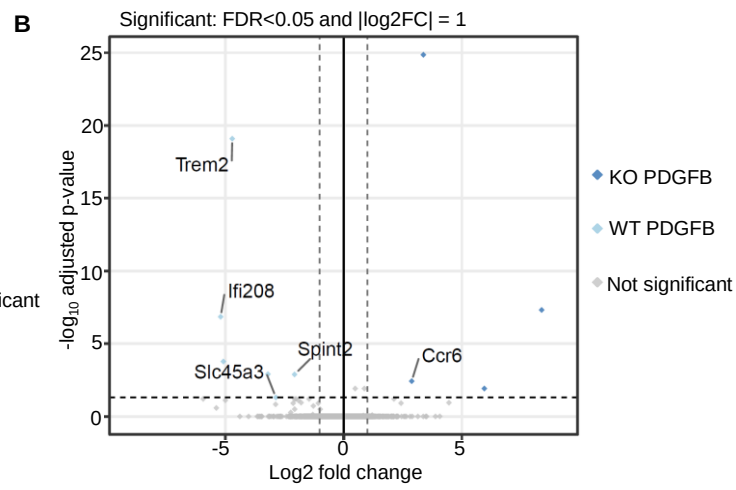

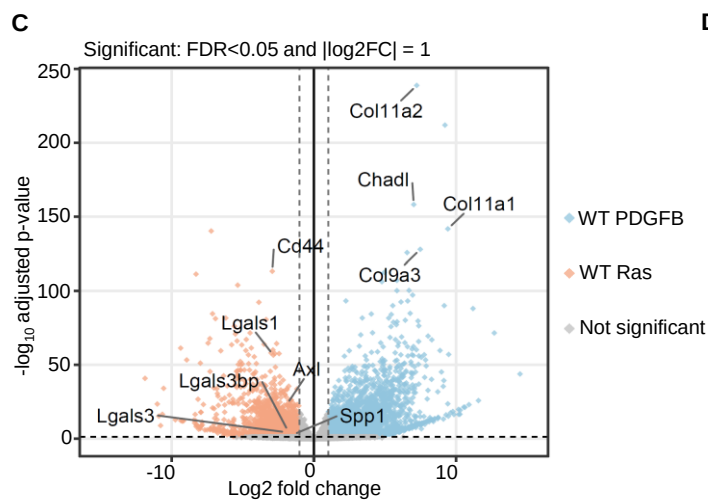

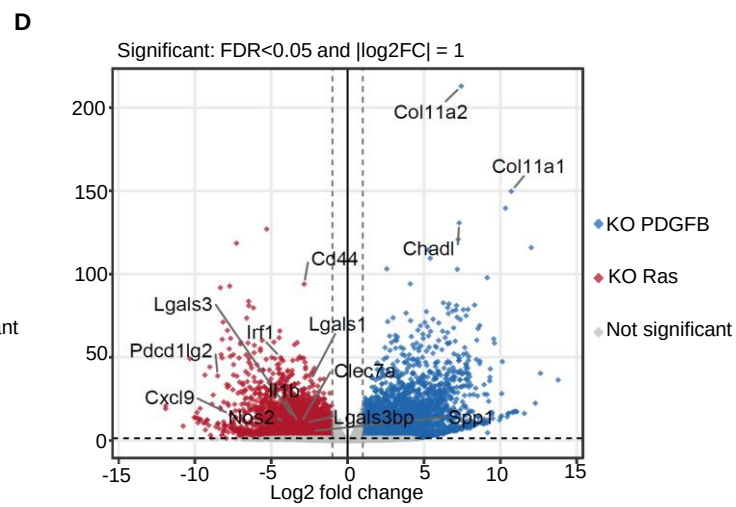

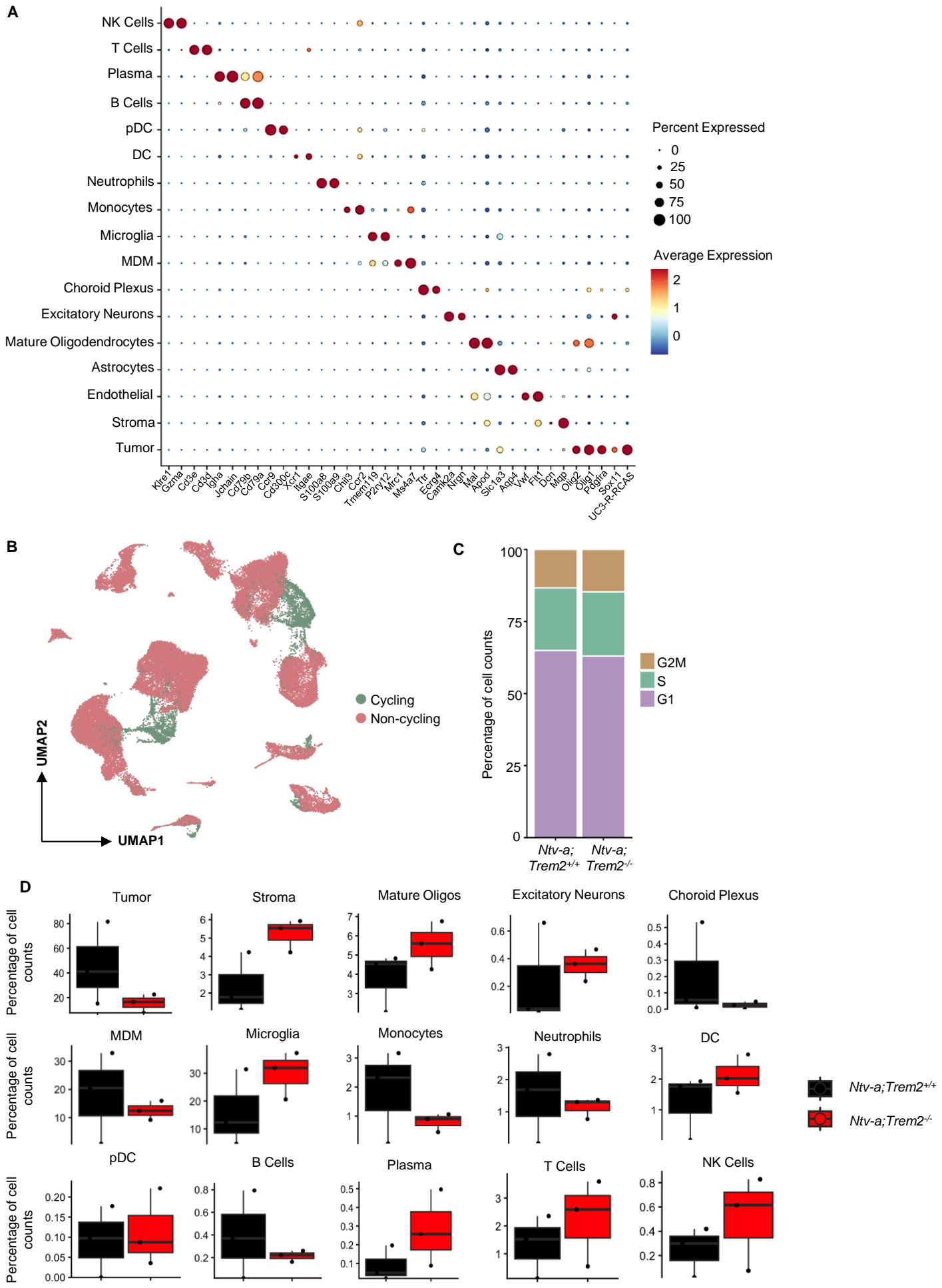

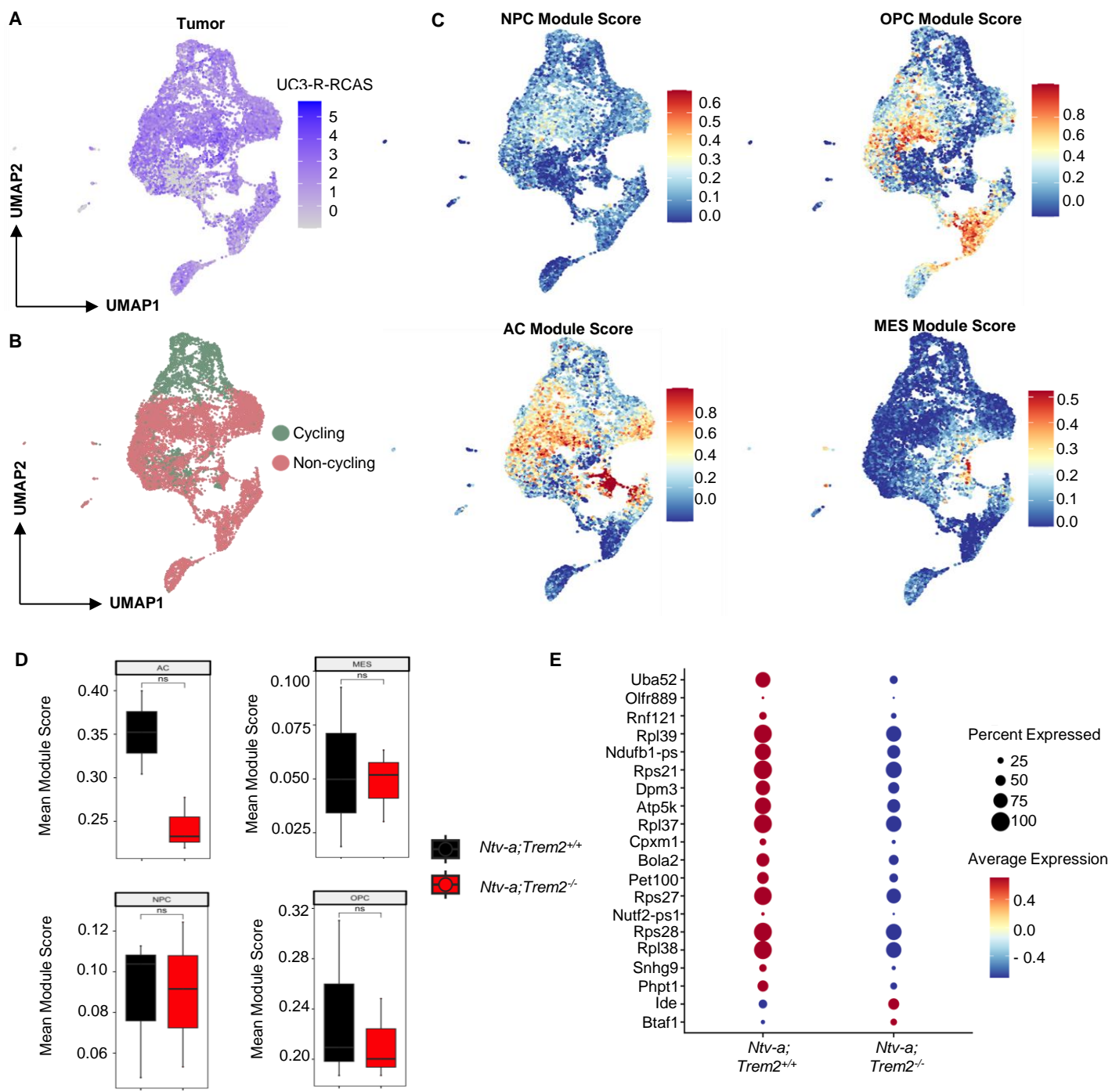

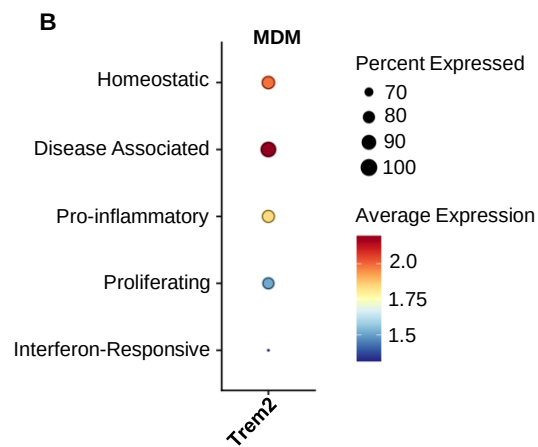

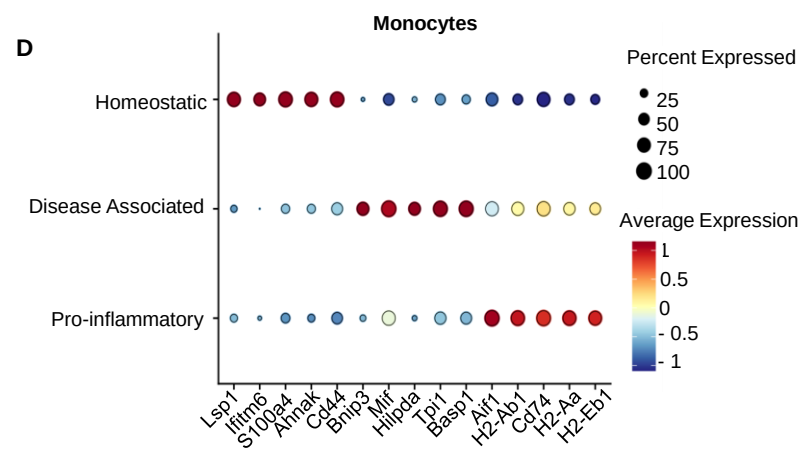

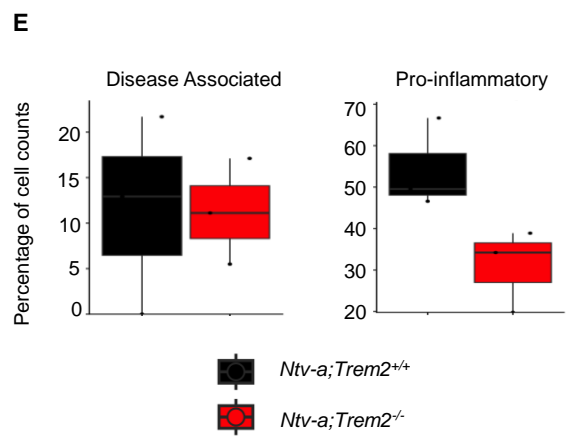

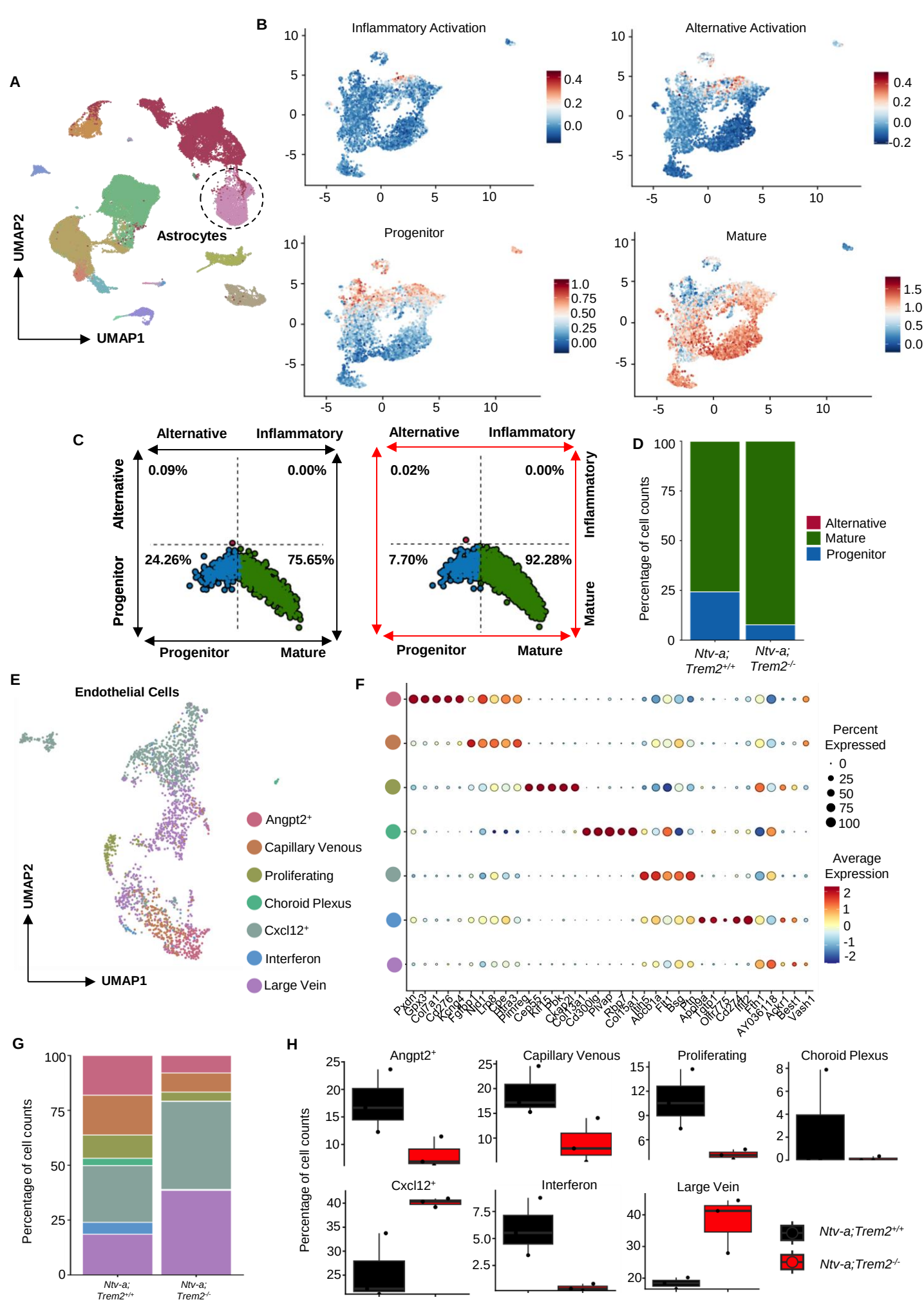

**A**

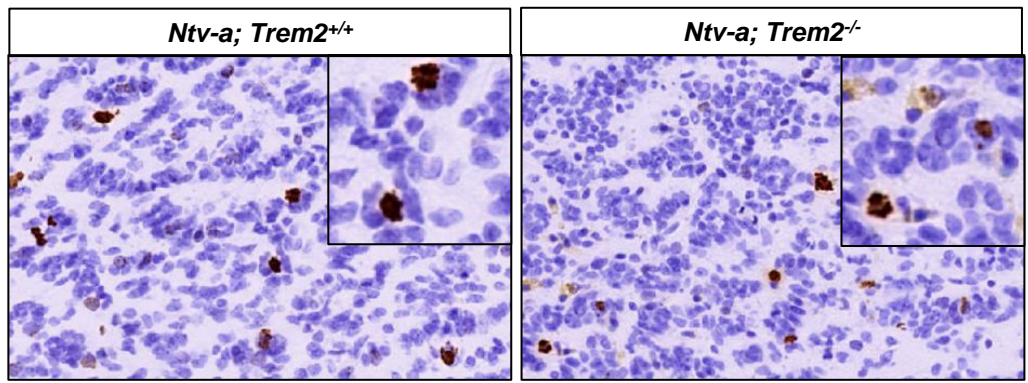

**B**

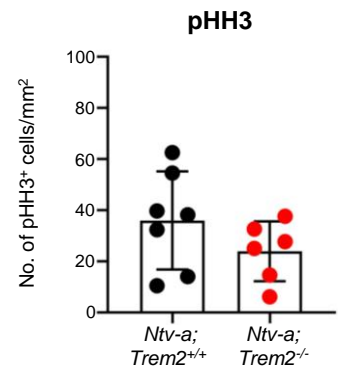

**C**

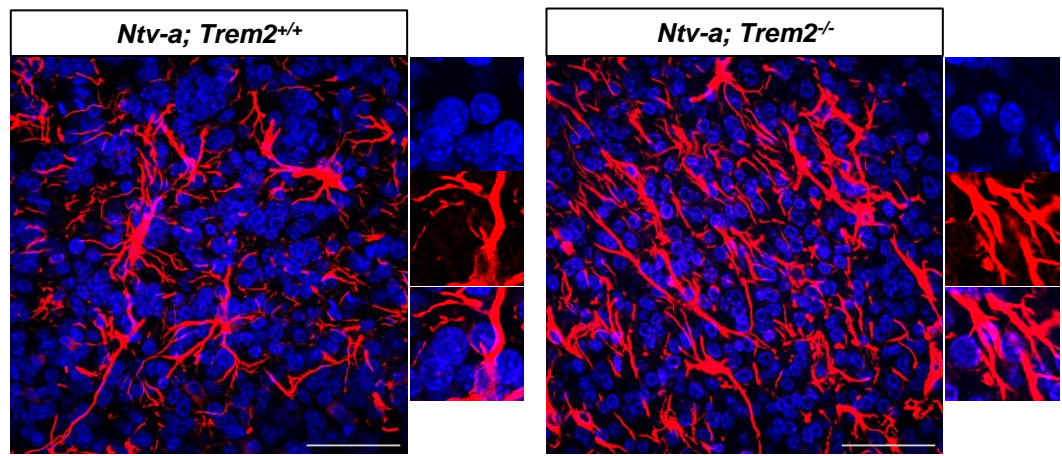

D

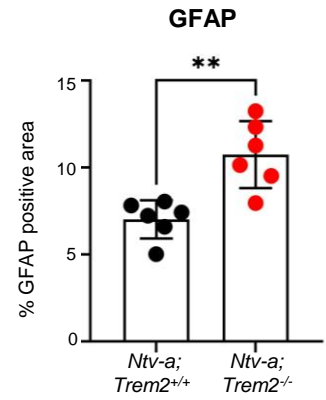

**E**

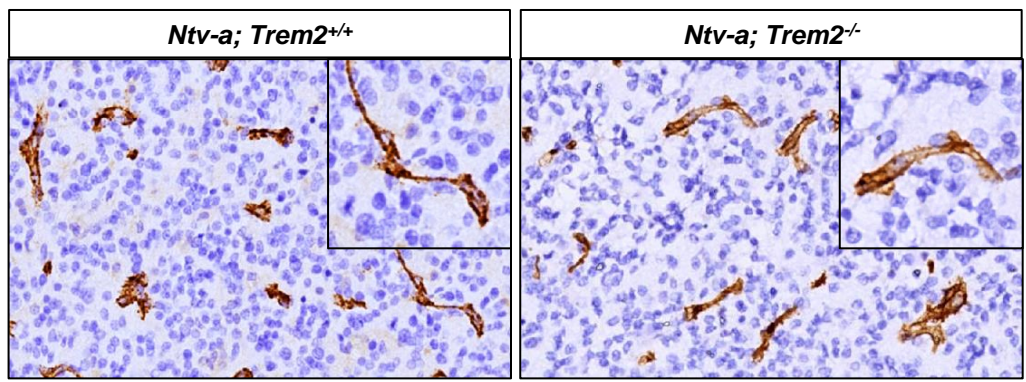

**F**

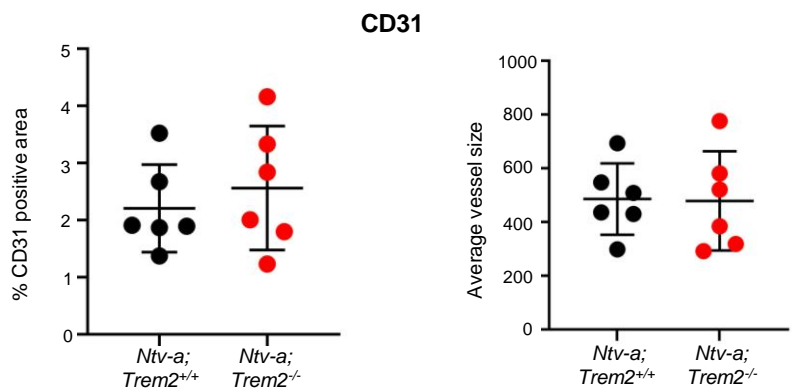

**A**

Collect brains from naïve *Trem2<sup>fllox/fllox</sup>* and *CX3CR1<sup>GFP/+</sup>; Trem2<sup>fllox/fllox</sup>* mice after tamoxifen administration

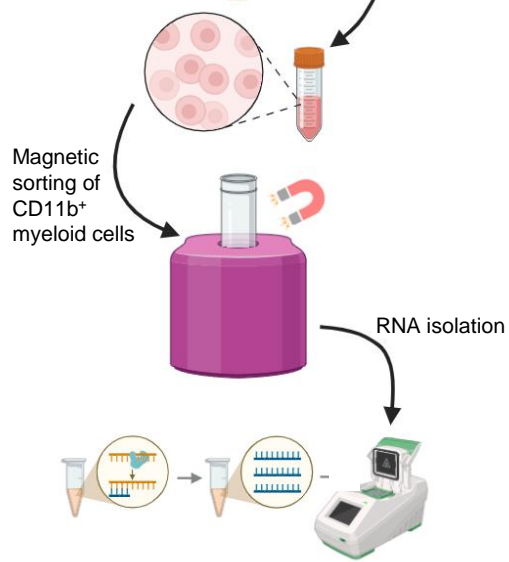

Assess *Trem2* expression by rt-qPCR

**B**

### Validation of Trem2 conditional knockout

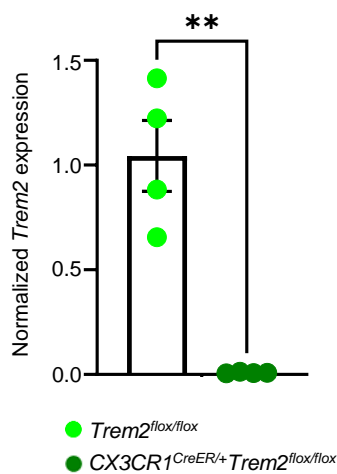
