## Supplemental Table 1 for "Genetic drivers shape TREM2-dependent myeloid responses and glioblastoma progression"

**Table 1: Antibodies used in this study.**

| **Name** | **Vendor** | **Catalogue** | **Dilution** | **Application** |
| --- | --- | --- | --- | --- |
| IBA1 | Wako | #019-19741 | 1:250 | IF, IHC |
| pHH3 | Cell Signaling | #9701 | 1:100 | IF, IHC |
| CD31 | Dianova | #Dia-310 | 1:50 | IHC |
| GFAP | Dako | *#*Z0334 | 1:500 | IF |
| TREM2 | Proteintech | #68723 | 1:100 | IF |
| Species-specific secondary antibodies | Jackson ImmunoResearch | N/A | 1:500 | IF |
| Goat anti-rabbit IgG, Alexa Fluor 488 | Invitrogen | #A-11008 | 1:500 | IF |
| CD45 PE/CF594 | BD Biosciences | #562420 | 1:1000 | FC |
| CD11b PE/Cy5 | Tonbo | #55-0112-U100 | 1:1000 | FC |
| CD3 PE/Fire700 | Biolegend | #100272 | 1:100 | FC |
| CD4 BV510 | Biolegend | #100449 | 1:50 | FC |
| CD8α BV785 | Biolegend | #100750 | 1:200 | FC |
| Ly6C Pacific Blue | Biolegend | #128014 | 1:100 | FC |
| Ly6G APC | Biolegend | #127614 | 1:100 | FC |
| B220 SparkNIR685 | Biolegend | #103268 | 1:100 | FC |
| NK1.1 APC/Fire810 | Biolegend | #156519 | 1:100 | FC |
| CD11b PerCP-Cy5.5 | BD Biosciences | #550993 | 1:100 | FC |
| CD101 APC | Invitrogen | #17-1011-80 | 1:100 | FC |
| CD4 APC-Cy7 | Biolegend | #100526 | 1:100 | FC |
| B220 BV605 | Biolegend | #103243 | 1:100 | FC |
| NK1.1 BV711 | BD Biosciences | #740663 | 1:100 | FC |
| PD-1 BV785 | Biolegend | #135225 | 1:100 | FC |
| CD45 BV510 | Biolegend | #103137 | 1:100 | FC |
| CD8 PE-Cy5 | BD Biosciences | #553034 | 1:100 | FC |
| CD3 PE/D | Biolegend | #100348 | 1:100 | FC |
| Tim-3 PE/Cy7 | Invitrogen | #25-5870-82 | 1:100 | FC |
| GrzmB PE (IC) | Invitrogen | #12-8899-41 | 1:50 | FC |
| FoxP3 FITC (IC) | Invitrogen | #11-5773-82 | 1:50 | FC |
| Ki67 eF506 (IC) | Invitrogen | #69-5698-80 | 1:50 | FC |
| CD45 FITC | Invitrogen | #11-5773-82 | 1:50 | FC |
| CD11c APC | BD Biosciences | #550261 | 1:100 | FC |
| MHC II A700 | Invitrogen | #56-5321-80 | 1:100 | FC |
| CD103 BUV395 | BD Biosciences | #748253 | 1:100 | FC |
| CD24 BUV496 | BD Biosciences | #612953 | 1:200 | FC |
| PD-L1 BV605 | BD Biosciences | #745135 | 1:100 | FC |
| CX3CR1 BV650 | Biolegend | #149033 | 1:100 | FC |
| F4/80 BV711 | Biolegend | #123147 | 1:100 | FC |
| Ly6G V450 | BD Biosciences | #560603 | 1:100 | FC |
| CXCR2 PE | Biolegend | #149609 | 1:100 | FC |
| CD49d PE/D | Biolegend | #103625 | 1:100 | FC |
| Ly6C PE/Cy7 | BD Biosciences | #560593 | 1:200 | FC |
| Zombie UV | Biolegend | #423107 | 1:1000 | FC |

Antibodies used in this study. IF=immunofluorescence, IHC=immunohistochemistry, FC=flow cytometry, IC=intracellular staining.
